# Iron uptake mediated by TFRC and secretion of transferrin types stimulate thermogenic activation in human adipocytes

**DOI:** 10.1101/2025.11.24.690185

**Authors:** Rahaf Alrifai, Mizuki Seo, Gyath Karadsheh, Fachrur Rizal Mahendra, Máté Á Demény, Ferenc Győry, János András Mótyán, László Fésüs, Endre Kristóf, Rini Arianti

**Affiliations:** Laboratory of Cell Biochemistry, Department of Biochemistry and Molecular Biology, Faculty of Medicine, University of Debrecen, Debrecen 4032, Hungary; Doctoral School of Molecular Cellular and Immune Biology, University of Debrecen, Debrecen 4032, Hungary; Department of Biochemistry, Faculty of Mathematics and Natural Sciences, IPB University, Dramaga Campus, Bogor 16680, Indonesia; Bioinformatics Research Center, Indonesian Institute of Bioinformatics (INBIO Indonesia), Malang, 65145, Indonesia; Department of Medical Chemistry, Faculty of Medicine, University of Debrecen, Debrecen 4032, Hungary; Department of Surgery, Faculty of Medicine, University of Debrecen, Debrecen 4032, Hungary; Laboratory of Retroviral Biochemistry, Department of Biochemistry and Molecular Biology, Faculty of Medicine, University of Debrecen, Debrecen 4032, Hungary

## Abstract

Adrenergic-driven thermogenic activation of brown adipose tissue requires high amounts of nutrients including iron supporting mitochondrial biogenesis. This was governed by rapid gene expression changes in human cervical-derived brown adipocytes. Transferrin receptor 1 (TFRC) was strongly upregulated in response to dibutyryl-cAMP. Pharmacological inhibition and siRNA-mediated knockdown of TFRC during adrenergic stimulation decreased intracellular iron content and prevented elevation of oxygen consumption and induction of thermogenic markers. Deferoxamine-mediated iron chelation also showed comparable effects. Contrarily, the expression of ferroportin iron exporter was suppressed during activation, however, its pharmacological inhibition did not further increase thermogenesis. Brown adipocytes constitutively expressed and secreted high amounts of transferrin while the released melanotransferrin at activated condition was predicted to further augment iron influx stimulating heat production.

## Introduction

Adipose tissue is recognized not only for its role in regulating energy balance and glucose homeostasis, but also for its involvement in diverse pathophysiological processes mainly with regard to metabolic diseases such as obesity and diabetes mellitus [Shamsi *et al*., 2021, *Nat. Rev. Endocrinol.*]. Three distinct types of adipocytes have been classified in mammals according to their thermogenic potential: non-thermogenic white adipocytes and two heat producing subtypes including classical brown and inducible brite (brown-in-white) or beige adipocytes [Sakers *et al*., 2022, *Cell*]. White adipose tissue (WAT), which is the most abundant form of adipose tissue, plays a critical role in nutrient homeostasis by storing energy in the form of triacylglycerols produced by the esterification of fatty acids with glycerol. Unlike WAT, brown adipose tissue (BAT) dissipates energy as heat mainly via the activity of uncoupling protein-1 (UCP1) which facilitates proton leak across the inner mitochondrial membrane [Rosen & Spiegelman, 2014, *Cell*].

In humans, BAT has been considered only to be found in newborns for a long period of time. However, nearly two decades ago, independent studies demonstrated the presence of BAT in the deep cervical (DeepC) depot of adults [Virtanen *et al*., 2009, *NEJM*; Cypess *et al*., 2009, *NEJM*]. Using a refined technique of positron emission tomography/computed tomography (PET/CT), Leitner *et al*. (2017) observed that brown or brownable adipose tissue can be found in DeepC, supraclavicular, axillary, mediastinal, paraspinal, and abdominal regions. These areas are rich in both classical brown and beige adipocytes distinguished by multilocular lipid droplets and a high density of mitochondria possessing dense cristae [Tóth *et al*., 2020, *Cells*; Jespersen *et al*., 2013, *Cell Metab.*]. Efficient heat production requires activation primarily by the sympathetic nervous system (SNS) that densely innervates BAT. The released norepinephrine binds to β-adrenergic receptors, which induce a signaling cascade leading to adenylyl cyclase activation, then the produced 3’,5’-cyclic adenosine monophosphate (cAMP) transmits the thermogenic signal mainly via Protein Kinase A [Cannon and Nedergaard, 2004, *Physiol Rev*]. This results in stimulation of lipolysis, fatty acid elongation, mitochondrial biogenesis, UCP1-dependent and independent thermogenesis, and characteristic gene expression changes [Kajimura *et al*., 2015, *Cell Metab*; Arianti *et al*., 2024, *Sci. Rep.*].

Thermogenically active adipocytes utilize high amounts of nutrients [Townsend and Tseng, 2014, *Trends Endocrinol. Metab.*] either for fueling catabolism or as structural components in various proteins. In association with its high metabolic activity, BAT accounts for approximately 5% of the basal metabolic rate in adult humans, equivalent to the oxidation of around 4 kg of fat per year [van Marken Lichtenbelt & Schrauwen, 2011, *Am. J. Physiol.-Regul., Integr. and Compar. Physiol.*]. Mitochondrial biogenesis and thermogenesis require iron as an essential cofactor for the synthesis of iron–sulfur clusters and heme groups incorporated into respiratory chain complexes [Read *et al*., 2021, *Redox Biol.*; Belot *et al*., 2024, *Liver Int.*]. In rodents, iron deficiency resulted in impaired adaptive thermogenic capacity of BAT and contributed to the development of obesity [Yook *et al*., 2021, *JBC*; Yook *et al*., 2021, *Journal of Nutrition*; Yook et *al.*, 2021, *PNAS*].

Intracellular iron homeostasis is mediated by the cell autonomous changes in the expression levels of iron transporter and storage proteins regulated primarily at post-transcriptional level by two cytosolic RNA-binding proteins, iron regulatory protein (IRP) 1 and IRP2, in response to iron availability. Transferrin receptor 1 (TFRC) is a homodimer glycoprotein composed of two 90 kDa subunits linked by disulfide bonds. It binds extracellular transferrin (TF) proteins loaded with ferric iron (Fe^3+^) in both lobes and mediates cellular iron uptake via receptor-mediated endocytosis. Within endosomes, Fe^3+^ is reduced to ferrous iron (Fe^2+^) and subsequently transported across the endosomal membrane into the cytoplasm. After iron release, the apo-TF– TFRC complex is recycled to the plasma membrane, where apo-TF dissociates at physiological pH, allowing the receptor to initiate another uptake cycle [Galy *et al*., 2024, *Nat. Rev. Mol. Cell Biol.*]. Melanotransferrin (MELTF) is a TF homologue, which can be either cell membrane bound or secreted, with a significant iron binding ability [Sekyere and Richardson, 2000, *FEBS Lett*]. Homeostatic iron regulator (encoded by *HFE*) can also bind to TFRC to regulate cellular iron uptake [Bennett *et al*., 2000, *Nature*]. Ferroportin, encoded by *SLC40A1*, transports iron from the inside to the outside of the cell, playing a key role in maintaining not only intracellular but also systemic iron homeostasis [Mitchell *et al*., 2014, *Am J Physiol Cell Physiol.*].

In order to identify critical molecular regulators of thermogenesis activation in human adipocytes, adipose-derived stromal cells (ASCs) were isolated from paired DeepC and cervical subcutaneous (SubQ) adipose tissue biopsies. Adipocytes differentiated from DeepC ASCs displayed high, while the SubQ-derived ones had limited but significant thermogenesis potential, respectively [Tóth *et al*., 2020, *Cells*]. When a global transcriptomic profile analysis was performed, we found that adrenergic stimulation significantly increased the expression of *TFRC* and *MELTF*, downregulated *SLC40A1*, while the expression of *TF* remained constantly high [Arianti *et al*., 2024, *Sci. Rep.* available at Sequence Read Archive database under accession number PRJNA1093362]. Previous studies concluded that iron availability and its TFRC-mediated influx are critical for the thermogenic competency of murine brown and beige adipocytes [Yook *et al*., 2021, *JBC*; Yook *et al*., 2021, *J. Nutr.*; Li *et al*., 2020, *Adv. Sci*.; Qiu *et al*., 2020, *Front. Cell. Dev. Biol.*]. However, data is limited about the requirement and the mechanism of facilitated uptake of iron in response to adrenergic-driven thermogenesis activation, especially in human cell models. We found that the prevention of activation induced iron influx by a chelator or pharmacological inhibition and silencing of TFRC significantly repressed the thermogenic response of both SubQ and DeepC-derived adipocytes. Furthermore, we observed constant high expression and secretion of TF and a strong upregulation of MELTF by cervical-derived adipocytes suggesting that the release of iron binding proteins by thermogenic adipocytes can stimulate iron accumulation and heat production during adrenergic-driven activation.

## Materials and methods

### Materials

All chemicals were obtained from Sigma-Aldrich (Munich, Germany) unless otherwise specified.

### Methods

#### Ethics Statements and obtained tissue samples

The collection of tissue samples was approved by the Medical Research Council of Hungary (20571-2/2017/EKU) in accordance with the EU Member States’ Directive 2004/23/EC regarding presumed consent for tissue collection. All experiments complied with the guidelines of the Helsinki Declaration. Informed consents were obtained from all participants prior to the surgical procedure. Paired biopsy samples from SubQ and DeepC adipose tissues were obtained during thyroid surgery to diminish the inter-individual variability. Patients with pre-existing diabetes mellitus, malignant tumors, or abnormal thyroid hormone levels at the time of surgery were excluded from the study. Human ASCs were isolated from the stromal-vascular fraction (SVF) of SubQ and DeepC adipose tissue biopsies, as previously described [Arianti *et al*., 2024, *Sci. Rep.*; Tóth *et al*., 2020, *Cells*]. Briefly, after the connective tissues were removed, the tissues were cut into small pieces, digested in PBS containing 120 U/mL collagenase (cat#c0773), and incubated at 37°C. Next, cell suspension was centrifuged at 1300 rpm for 7 minutes and SVF pellets were resuspended in Dulbecco’s Modified Eagle’s Media/Nutrient F-12 Ham (DMEM-F12, cat#T2252) medium supplemented with 10% Fetal Bovine Serum (Thermo Fisher Scientific, Waltham, MA, USA, cat#A5256701), 100 U/ml penicillin-streptomycin (Roche, Basel, Switzerland, cat#11074440001), 33 μM biotin (cat#B4639), and 17 μM pantothenic acid (cat#P5155). The absence of Mycoplasma was verified utilizing a polymerase chain reaction (PCR) Mycoplasma Test Kit I/C (Promocell, Heidelberg, Germany, cat#PK-CA91).

#### Differentiation and treatment of human ASCs

Primary human SubQ and DeepC-derived adipocytes were differentiated from SVF according to a described adipogenic differentiation protocol to obtain cells with limited and high thermogenic potential, respectively [Arianti *et al*., 2024, *Sci. Rep.*]. Differentiation was induced using serum-free DMEM-F12 medium supplemented with 33 μM biotin, 17 μM pantothenic acid, 100 U/mL penicillin/streptomycin, 10 μg/mL human apo-TF (cat#T2252), 20 nM human insulin (cat#I9278), 200 pM triiodothyronine (cat#T6397), 100 nM hydrocortisone (cat#H0888), 2 μM rosiglitazone (Cayman Chemicals, Ann Arbor, MI, USA, cat#71740), 25 nM dexamethasone (cat#D1756), and 500 μM 3-isobutyl-1-methylxanthine (IBMX, cat#I5879). After 3 days, rosiglitazone, dexamethasone, and IBMX were omitted from the differentiation medium. After 14 days of differentiation, mature adipocytes were treated with 500 μM dibutyryl (db)-cAMP (cat#D0260) for 10 hours to activate them for thermogenesis, in the presence or absence of 10 μM iron chelator desferoxamine (DFO, cat#D9533) [Kremer *et al*., 2021, *Nat. Comm*.], or 200 nM ferroportin antagonist (VIT-2763, Med Chem Express, Monmouth Junction, NJ, USA, cat# HY-112220), or 50 μM Ferristatin II (cat#NSC8679) TFRC inhibitor [Byrne *et al*., 2013, *Plos One*].

#### Small-interfering RNA (siRNA)-mediated knock-down of TFRC in human DeepC-derived adipocytes

Transfection and gene silencing experiments were performed using DharmaFECT1 transfection reagent (Dharmacon, Lafayette, CO, USA, cat#T-2001-03) at day 14 of differentiation by following the manufacturer’s instruction. Mature DeepC-derived adipocytes were incubated with mixture of DharmaFECT1 and 100 nM of TFRC-targeting siRNA (Dharmacon ON-TARGETplus SMARTpool Human TFRC siRNA, cat#L-003941-00-0005) or 100 nM non-targeting (scrambled) siRNA (Dharmacon plus non-targeting control pool, cat#D-001810-10-20) for 48 hrs at 37°C with 5% CO_2_. Following the incubation time, adipocytes were activated with 500 μM db-cAMP for 10 hours.

#### RNA isolation and RT-qPCR

Cells were collected and total RNA was isolated as described previously [Tóth *et al*., 2020, *Cells*]. The concentration and purity of the isolated RNA was checked by using Nanodrop 2000 Spectrophotometer (Thermo Fisher Scientific). RNA was diluted to 100 ng/µL for all samples and was reverse transcribed to cDNA by using reverse transcription kit (Thermo Fisher Scientific, cat#4368814) following the manufacturer’s instructions. Validated TaqMan assays used in qPCR were designed and supplied by Thermo Fisher Scientific as listed in **Supplementary Table 1**. qPCR was performed in Light Cycler 480 II (Roche). The following conditions were set to perform the reactions: initial denaturation step at 95 °C for 1 min followed by 40 cycles of 95 °C for 12 s and 60 °C for 30 s. Gene expression values were calculated by the comparative threshold cycle (Ct) method as described in the previous publication [Huang *et al*., 2023, *Plos Biology*]. Gene expressions were normalized to the geometric mean of *ACTB* and *GAPDH*. Normalized gene expression levels equal 2^-ΔCt^.

#### Immunostaining and fluorescent imaging

Differentiated and activated adipocytes were washed and fixed in 4% paraformaldehyde for 5 min followed by staining with anti-TFRC (1:100, Santa Cruz Biotechnology, Dallas, TX, USA, cat#K1821) primary antibody at room temperature overnight. Alexa 488 goat anti-mouse IgG (1:1000, Thermo Fisher Scientific, cat#1298479) was used as a secondary antibody. Antibodies were applied and additional washing steps between and after antibody usage were performed in the presence of 0.1% saponin in PBS for effective cell permeabilization. Finally, cells were stained with 20 mM Hoechst 33342 (1:10000, Thermo Fisher Scientific, cat#VL3148741) and AdipoRed (1:40, Lonza, Basel, Switzerland, cat#PT-7009) for 15 minutes. Confocal imaging was performed on a Leica (Wetzlar, Germany) TCS SP8 system coupled to a Leica DM6000 microscope using a Leica HC PL APO 63×/1.4 NA OIL CS2 objective. Laser power, detector gain, and offset were kept constant between samples, and the pinhole was maintained at 1 Airy unit. 1024x1024 scan format and 0.75x zoom factor were used to include more cells. Image acquisition was performed in bidirectional scanning mode at 100 Hz with PhaseX correction. Owing to the thickness of lipid droplet–filled adipocytes, images were acquired as Z-stacks with a step size of 0.4 µm. Fluorescence was recorded in three channels in sequential mode, and final images represent sum intensity projections generated in FIJI ImageJ software (National Institutes of Health, Bethesda, MD, USA). The Hoechst and Nile Red channels, respectively marking nuclei and lipid droplets, were overlaid on the immunofluorescent signal of the proteins of interest to aid spatial interpretation.

#### Immunoblotting and densitometry

Immunoblotting and densitometry were carried out as described previously [Szatmári-Tóth *et al*., 2020, *IJMS*]. Differentiated adipocytes were washed with ice-cold PBS, lysed by using the denaturing agent SDS, and then collected to be loaded onto a SDS polyacrylamide gel. Subsequently, the proteins were transferred onto PVDF Immobilon-P Transfer Membranes (Merck Millipore Ltd., Cork, Ireland, cat#IPVH00010), followed by a blocking step in Tris-buffered saline containing 0.05% Tween-20 (TBS-T) and 5% skimmed milk for 1 hour. Antibodies and working dilutions are listed in **Supplementary Table 2**. The expression of the visualized immunoreactive proteins were quantified by densitometry using the FIJI ImageJ software as previously described [Arianti *et al*., 2024, *Sci. Rep.*; Vinnai *et al*., 2023, *Front. Nutr.*].

#### Iron content measurement

Mature adipocytes which were treated with db-cAMP, DFO, VIT-2763, or combination of db- cAMP and DFO or VIT-2763 were homogenized in 50 µl of the iron assay buffer. Intracellular iron levels in TFRC-silenced adipocytes were also measured similarly. Total iron content (Fe^2+^ and Fe^3+^) levels were quantified by using a commercially available iron assay kit (cat#MAK025) according to the manufacturer’s protocols. Iron levels were quantified in the conditioned culture medium following ferristatin II treatment. To determine iron influx, the iron concentration in the conditioned cultured medium was measured and compared with that of the unconditioned cultured medium. Iron influx was calculated as the difference between the unconditioned and conditioned medium, where a lower iron concentration in the cultured medium indicated cellular iron uptake. The quantified iron levels were normalized to cellular protein contents which were measured by using Pierce™ BCA protein assay kit (Thermo Fisher Scientific, cat#23225).

#### Extracellular flux assay

Oxygen consumption rate (OCR) of adipocytes was measured using an XF96 oxymeter (Seahorse Biosciences, North Billerica, MA, USA) as described previously [Arianti *et al*., 2023, *JNB*]. The OCR was recorded at baseline and after the injection of db-cAMP, DFO, VIT-2763, ferristatin II, or combination of db-cAMP and one of the inhibitors. For TFRC knock-down experiments, the extracellular flux assay was performed upon 48-hour silencing by TFRC siRNA. Proton leak respiration was monitored after injecting oligomycin (cat#495455) at 2 μM concentration. Cells received a single bolus of Antimycin A (cat#A8674) at 10 μM concentration for baseline correction (measuring non-mitochondrial respiration). The OCR was normalized to protein contents which were measured by using Pierce™ BCA protein assay kit.

#### Measurement of human TF and MELTF secretion

The conditioned media after the 10 hours activation were collected and stored in –70 °C prior the measurement. ELISA kits were used to measure TF (Thermo Fisher Scientific, cat#EHTF) and MELTF (Novus Biologicals, Centennial CO, USA, cat#NBP3-39829) secretion. The concentration of both proteins in the conditioned media was calculated in accordance with the supplier’s instructions. The concentration of the secreted proteins was normalized to the protein contents of the cells per sample.

#### Interactome analysis and visualization

Protein-protein interactions among TFRC, MELTF, TF, and ferroportin was analyzed by using STRING (https://string-db.org) [Szklarczyk *et al*., 2023, NAR] and visualized by using Gephi 0.9 (The Gephi Consortium, Paris, France).

#### Single nuclei (sn) RNA-sequencing (RNA-seq) analysis

Available snRNA-seq dataset was explored by using public webtool (https://batnetwork.org/) as described by Sun *et al*., (2020).

#### Structural modeling and protein-protein interaction analysis

The interaction between MELTF and TFRC was predicted using AlphaFold2 and AlphaFold2- Multimer implemented through ColabFold v1.5.5 (https://colab.research.google.com/github/sokrypton/ColabFold/blob/main/AlphaFold2.ipynb) [Mirdita *et al*., 2022, *Nat. Methods*; Ibrahim *et al*., 2023, *Plos Biol.*]. Protein inputs included the TFRC dimer (PDB: 1SUV, chains A–B) and the MELTF monomer (PDB: 6XR0, chain C) [Hayashi *et al*., 2021, *Sci. Rep.*; Cheng *et al*., 2004, *Cell*]. The TFRC–TF complex structure was provided as a template to guide model refinement. Multiple sequence alignments were generated with mmseqs2_uniref_env, and structural templates were identified using HHsearch. The five top-ranked models were refined with the AMBER force field and evaluated using predicted Local Distance Difference Test (pLDDT), Predicted Aligned Error (PAE), and interface predicted Template Modeling (ipTM) scores [Wróblewski & Kmiecik, 2024, *Computat. Structur. Biotech. J.*]. Interfacial residues were identified with InterProSurf (https://curie.utmb.edu/usercomplex.html) [Negi *et al*., 2007, *Bioinformatics*] by using the TFRC–TF reference structure and the predicted TFRC–MELTF complex. Identified residues were used to define docking restraints. Docking simulations were performed using HADDOCK 2.4 [Honorato *et al*., 2024, *Nat. Protoc.*]. The docking protocol included rigid-body energy minimization (1,000 models), semi-flexible simulated annealing refinement (200 models), and explicit solvent refinement (200 models). Docking results were evaluated by HADDOCK score, Z-score, RMSD, and buried surface area (BSA). Two-dimensional interaction fingerprints were generated with PDBsum to identify hydrogen bonds, van der Waals, disulfide, and electrostatic interactions [Laskowski *et al*., 2018, *Prot. Sci*]. Three-dimensional conformations and molecular surfaces were visualized using PyMOL v3.1.6.1 (Schrödinger Inc, New York City, NY, USA).

#### Data visualization and statistical analysis

The data are expressed as the mean ± standard deviation (SD). The normality of data distribution was assessed by the Shapiro–Wilk test. Group comparisons were conducted using one-way analysis of variance (ANOVA), followed by Tukey’s *post hoc* test. All statistical analyses and data visualizations were generated using GraphPad Prism, version 8 (GraphPad Software, San Diego, CA, USA).

## Results

### Adrenergic stimulation induced the expression iron transport-related proteins while downregulated iron exporter in cervical area-derived adipocytes

We previously reported the global transcriptomic alterations in human SubQ and DeepC-derived adipocytes in response to db-cAMP treatment [Arianti *et al*., 2024, *Sci. Rep.*] and found that the expression of genes encoding iron transport-related proteins (**Fig. 1A**) were regulated upon thermogenic activation (**Fig. 1B**). *TFRC* was among the genes whose expressions were significantly induced by db-cAMP (**Fig. 1C**) and this result was confirmed by RT-qPCR (**Fig. 1D**). The expression of *TF* that encodes a key protein responsible for extracellular iron transport by adipocytes was also analyzed. Our RNA-seq data showed that it was constitutively highly expressed and not influenced by adrenergic-driven activation in either SubQ or DeepC-derived adipocytes (**Fig. 1E**), and RT-qPCR verified this expression pattern (**Fig. 1F**). In contrast, the expression of iron exporter ferroportin (encoded by *SLC40A1*) was suppressed by db-cAMP in both SubQ and DeepC-derived adipocytes (**Fig. 1G**). The mRNA expression data detected by RT-qPCR validated the similar expression pattern of ferroportin (**Fig. 1H**). Adrenergic stimulation driven by db-cAMP also induced the mRNA expression (**Fig. 1I and J**) of *MELTF*, especially in the thermogenically prone DeepC-derived adipocytes, which to date has not been recognized as an expressed iron transport-related protein by adipocytes. Our data suggest that thermogenically activated human adipocytes maintain sufficient intracellular iron levels by upregulating the expression of proteins required for iron uptake meanwhile downregulating ferroportin that is responsible for iron efflux.

**Figure 1.**
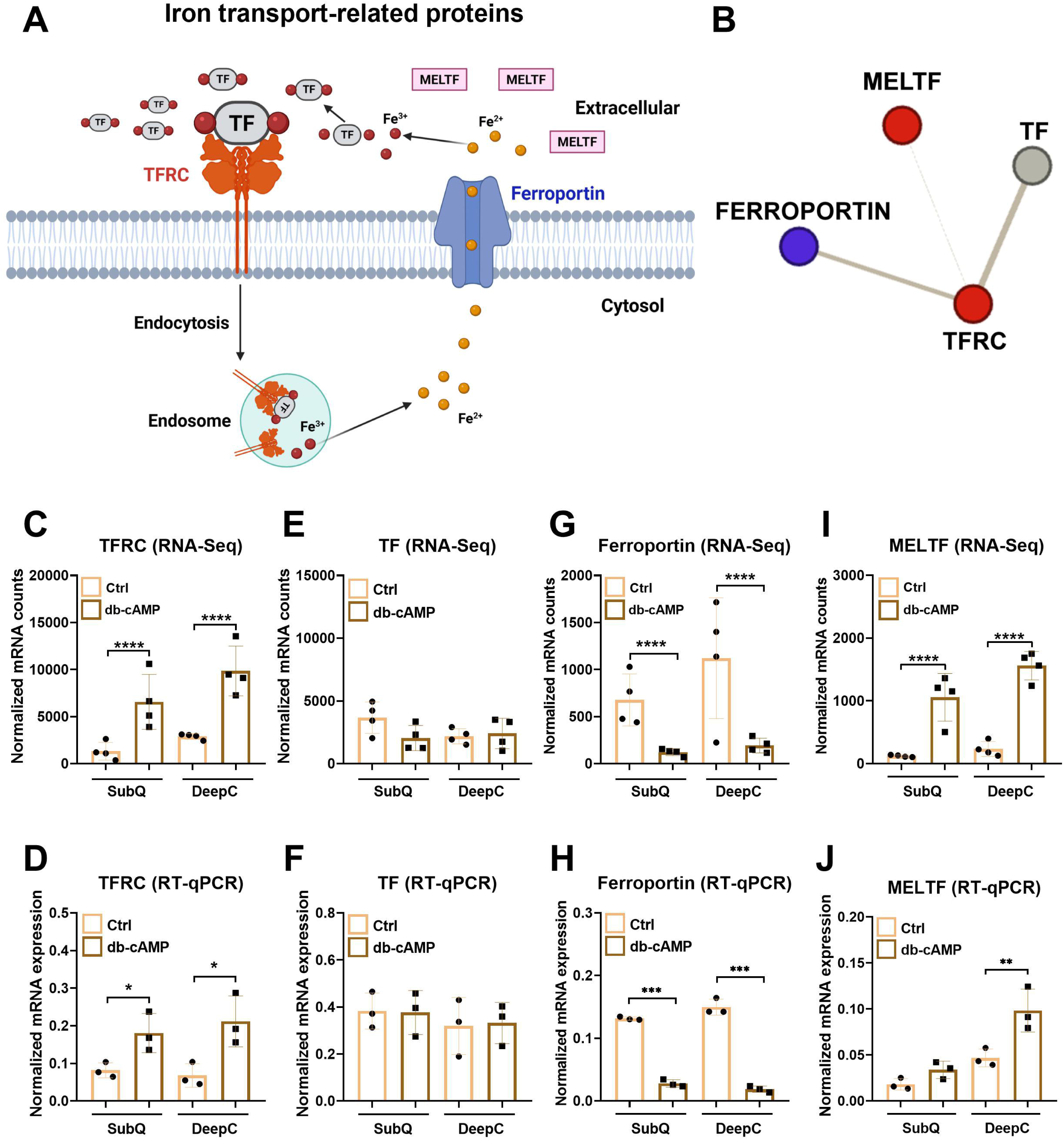
The effect of dibutyryl (db)-cAMP-driven stimulation of thermogenesis on the expression of iron transport-related proteins in human *ex vivo* differentiated subcutaneous (SubQ) and deep cervical (DeepC)-derived adipocytes. (A) Schematic figure displaying the iron transport proteins. Figure was created by Biorender. (B) Protein-protein interactions between transferrin receptor (TFRC), transferrin (TF), melanotransferrin (MELTF), and ferroportin based on STRING (www.string-db.org) analysis was generated by Gephi 0.9. Red: the mRNA expression was induced by db-cAMP in both SubQ and DeepC-derived adipocytes, blue: the mRNA expression was suppressed by db-cAMP in both SubQ and DeepC-derived adipocytes, and gray: no change in expression [see Supplementary Table 3-6 of Arianti *et al*., 2024, *Sci. Rep.*]). (C,E,G,I) Normalized mRNA counts of *TFRC* (C), *TF* (E), *SLC40A1/ferroportin* (G), and *MELTF* (I) based on RNA-sequencing data, n=4. The RNA-sequencing datasets generated and analyzed for this study can be found in the Sequence Read Archive (SRA) database [https://www.ncbi.nlm.nih.gov/sra] under accession number PRJNA1093362 [Arianti *et al*., 2024, *Sci. Rep.*]. Statistical analysis of FASTQ files aligned to BAM were performed by DESeq2. ****p<0.0001. (D,F,H,J) mRNA expression of *TFRC* (D), *TF* (F), *ferroportin* (H), and *MELTF* (J) in control (Ctrl) and activated adipocytes detected by RT-qPCR. n=3, statistical analysis was performed by one-way ANOVA followed by Tukey’s *post hoc* test, *p<0.05, **p<0.01, ***<0.001.

### DFO-mediated iron depletion reduced oxygen consumption and expression of mitochondrial complex subunits and thermogenic genes in human adrenergic activated adipocytes

Primarily, we aimed to investigate whether restriction of surplus iron availability prevents effective thermogenesis activation. We found that the intracellular iron contents were increased by db-cAMP in both SubQ and DeepC-derived adipocytes and this elevation was impaired by DFO (**Fig. 2A**). Next, we investigated the effect of DFO-mediated iron chelation on the oxygen consumption and found that stimulated maximal and proton leak respiration, which is associated with UCP1-dependent heat generation, were decreased during adrenergic stimulation in both SubQ (**Fig. 2B**) and DeepC-derived (**Fig. 2C**) adipocytes. We also found that DFO prevented the upregulation of mitochondrial complex I, II, and IV subunits during thermogenic activation by db-cAMP only in DeepC-derived adipocytes, except for the complex I subunit (**Fig. 2D**).

**Figure 2.**
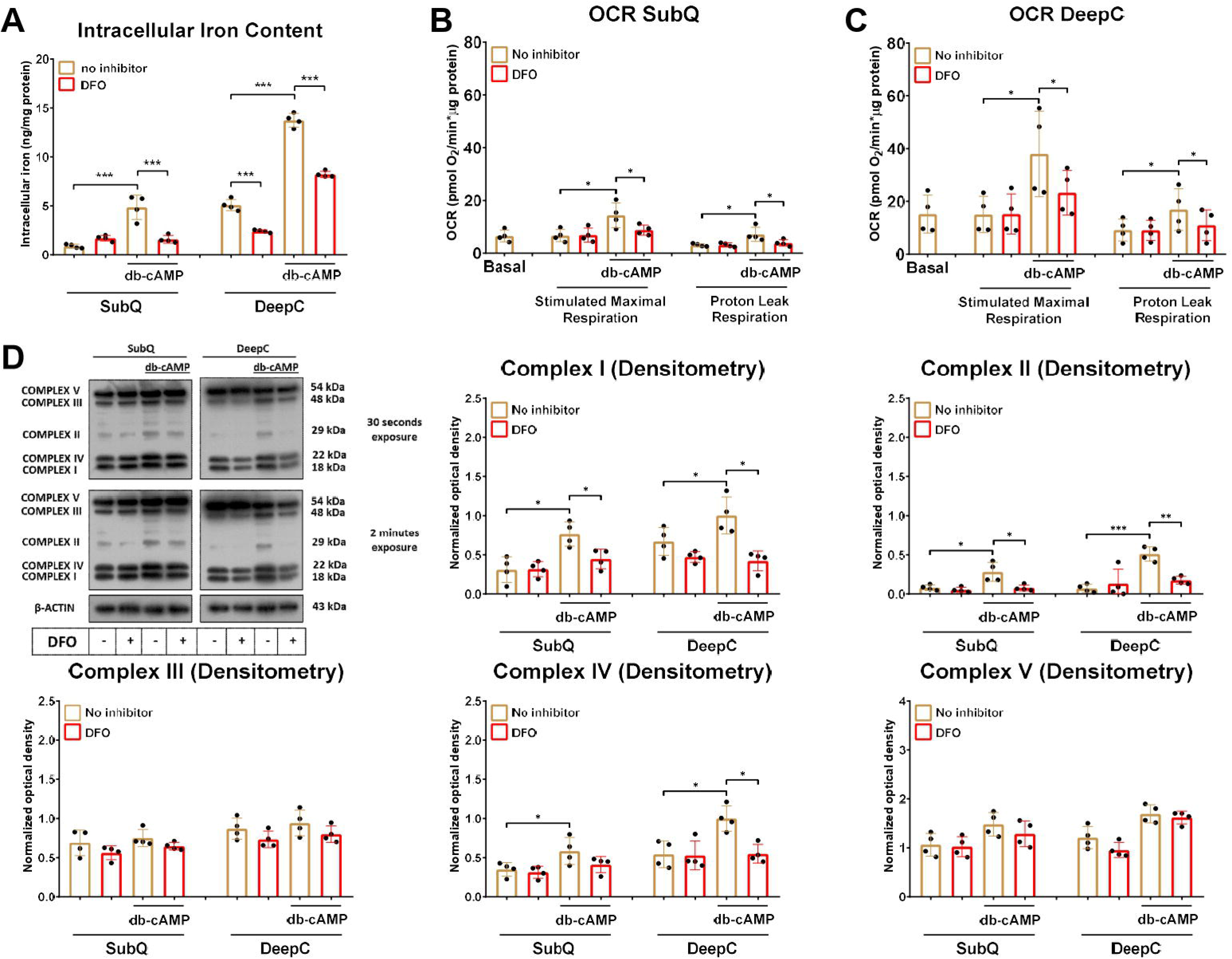
The effect of iron chelator deferoxamine (DFO) on oxygen consumption and protein expression of mitochondrial complex subunits in human *ex vivo* differentiated subcutaneous (SubQ) and deep cervical (DeepC)-derived adipocytes. SubQ and DeepC- derived adipocytes were treated with 500 µM dibutyryl (db)-cAMP, 10 µM DFO, or combination of the two compounds for 10 hours. (A) Intracellular iron content in adipocytes upon DFO treatment during adrenergic stimulation. (B-C) Basal, db-cAMP stimulated maximal, and proton leak oxygen consumption rate (OCR) in SubQ (B) and DeepC-derived (C) adipocytes were quantified by Seahorse extracellular flux analysis. (D) Protein expression of mitochondrial complex subunits detected by immunoblotting. The original uncropped images of the full-length blots are displayed in **Supplementary Fig. 3.** n=4, statistical analysis was performed by one-way ANOVA followed by Tukey’s *post hoc* test, *p<0.05 and ***p<0.001.

After observing the effect of DFO-mediated iron chelation on the oxygen consumption and heat generation, we were curious whether the expression of thermogenic genes was also affected. The db-cAMP-stimulated elevation of UCP1 and PGC1a was hampered by DFO at both mRNA and protein levels only in DeepC-derived adipocytes (**Fig. 3A, B**). We also observed that iron depletion decreased the protein expression of PGC1a in DeepC-derived adipocytes at unstimulated condition (**Fig. 3B**), likely due to impaired mitochondrial biogenesis which requires high amounts of iron. In addition, we also investigated the expression of *DIO2*, *CITED1*, and *PM20D1* and found that their adrenergic-driven upregulation was prevented in the presence of DFO (**Fig. 3C**). These data suggest that iron depletion inhibits the db-cAMP-stimulated elevation of intracellular iron content resulting in hampered oxygen consumption, mitochondrial complex subunit I, II, and IV and thermogenic gene expression in the cervical-derived adipocytes.

**Figure 3.**
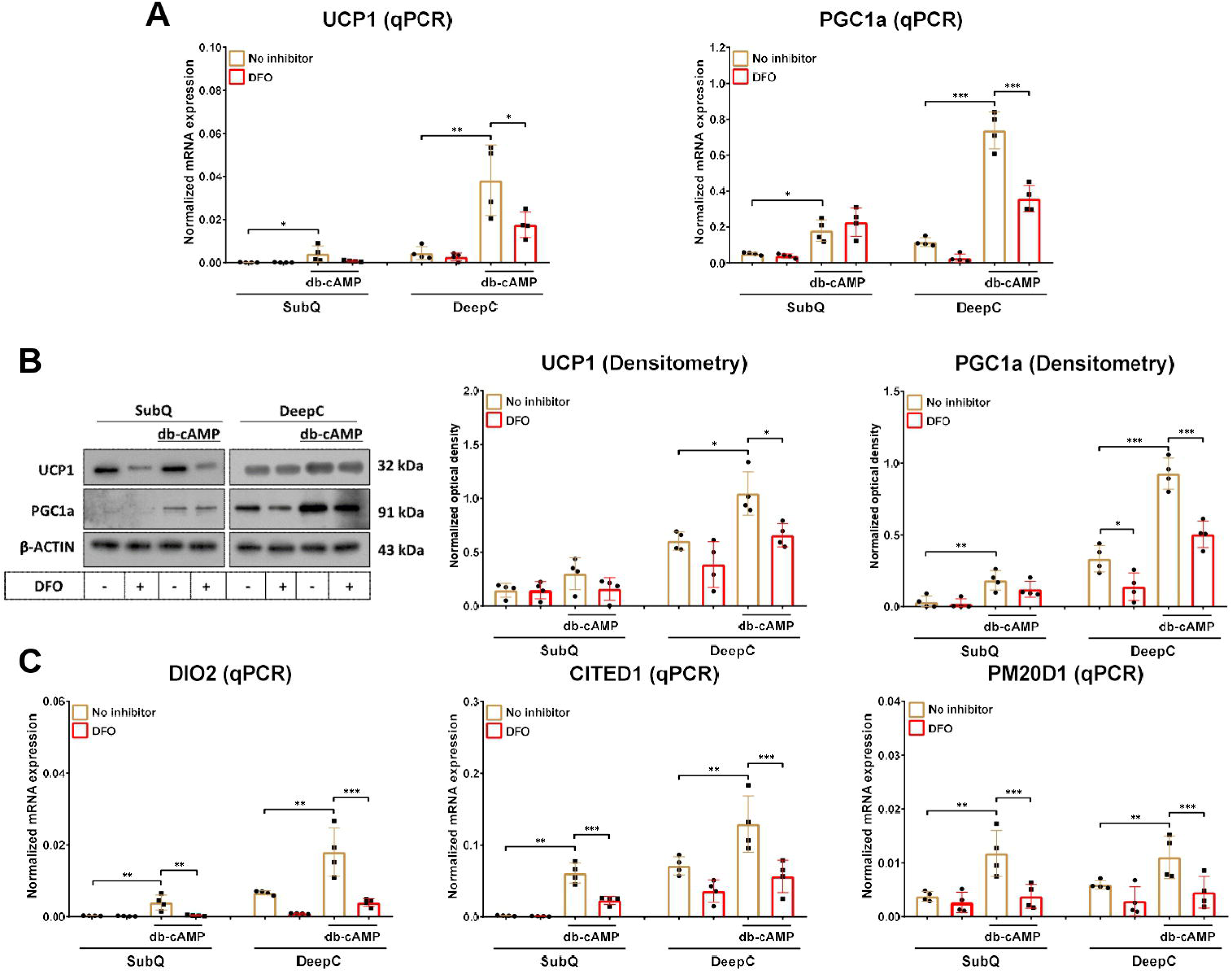
The effect of iron chelator deferoxamine (DFO) on the expression of thermogenic markers in human *ex vivo* differentiated subcutaneous (SubQ) and deep cervical (DeepC)- derived adipocytes. Adipocytes were differentiated and treated as in Figure 2. (A-B) mRNA (A) and protein (B) expression of UCP1 and PGC1a detected by RT-qPCR and immunoblotting, respectively. The original uncropped images of the full-length blots are displayed in **Supplementary Fig. 4.** (C) mRNA expression of *DIO2*, *CITED1*, and *PM20D1* detected by RT- qPCR. n=4, statistical analysis was performed by one-way ANOVA followed by Tukey’s *post hoc* test, *p<0.05, **p<0.01, and ***p<0.001.

### Inhibition of TFRC by ferristatin II decreased iron influx and the expression of mitochondrial respiratory chain proteins and thermogenic markers in human adrenergic activated adipocytes

As a next step, cervical-derived adipocytes were treated with ferristatin II to specifically inhibit TFRC by promoting its internalization and degradation via a lysosome-dependent pathway [Byrne *et al*., 2013, *Plos One*]. In parallel with mRNA expression shown in **Fig. 1C and D**, the protein expression of TFRC was elevated by adrenergic stimulation (**Fig. 4A, B**), primarily distributed in punctate structures in the cytoplasm (**Fig. 4A**), which was significantly decreased upon ferristatin II treatment at both control and db-cAMP stimulated conditions (**Fig. 4B**). Ferristatin II promotes clathrin-independent endocytosis of TFRC, therefore, instead of being recycled back to the plasma membrane, TFRC is redirected to the lysosome for degradation [Byrne *et al*., 2013, *Plos One*]. We also found that ferristatin II prevented the iron influx into DeepC-derived adipocytes during thermogenic activation (**Fig. 4C**). In parallel, the expression of mitochondrial complex subunits, which contain iron-sulfur clusters (complexes I, II, and III) or heme groups (complexes II, III and IV) in their structures [Read *et al*., 2021, *Redox Biol.*], was decreased by ferristatin II at both unstimulated and db-cAMP activated conditions in SubQ and DeepC-derived adipocytes (**Fig. 4D**). On the other hand, TFRC inhibition did not influence protein expression of complex V, which is composed of neither an iron-sulfur cluster nor a heme containing subunit in its structure [Read *et al*., 2021, *Redox Biol.*] (**Fig. 4D**). These data indicate that TFRC-mediated iron uptake is essential to maintain the high expression of iron-containing mitochondrial respiratory chain proteins during thermogenic activation.

**Figure 4.**
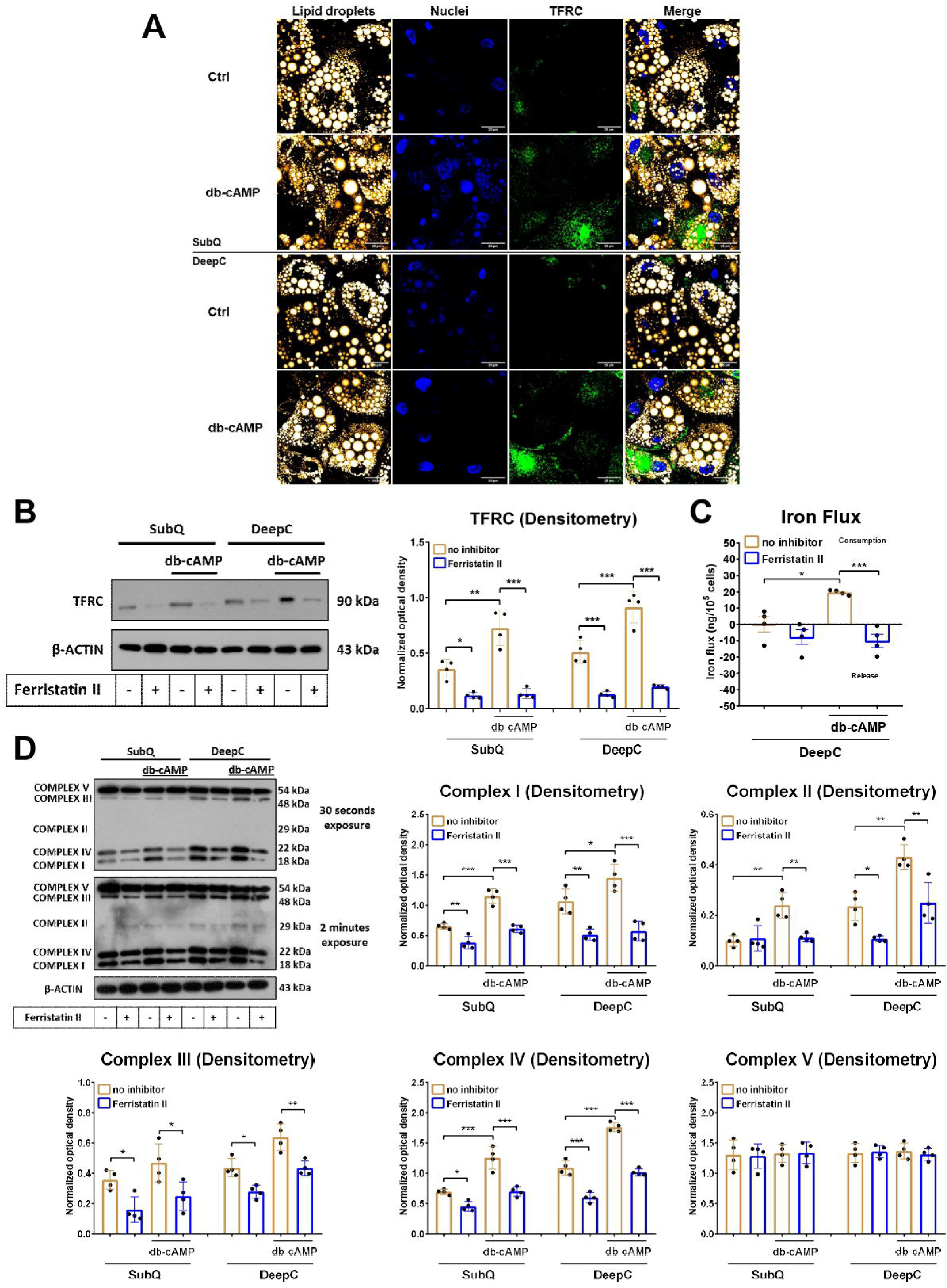
Inhibition of transferrin receptor (TFRC) by ferristatin II decreased the iron flux and mitochondrial complex subunits content in human *ex vivo* differentiated subcutaneous (SubQ) and deep cervical (DeepC)-derived adipocytes. SubQ and DeepC-derived adipocytes were treated with 500 µM dibutyryl (db)-cAMP, 50 µM ferristatin II, or combination of the two compounds for 10 hours. (A) Distribution of lipid droplets and TFRC in differentiated adipocytes. Bars represent 25 μm. (B) Protein expression of TFRC detected by immunoblotting. (C) Iron flux upon ferristatin II treatment during adrenergic stimulation in DeepC-derived adipocytes. (D) Protein expression of mitochondrial complex subunits detected by immunoblotting. The original uncropped images of the full-length blots are displayed in **Supplementary Fig. 5.** Statistical analysis was performed by one-way ANOVA followed by Tukey’s *post hoc* test, *p<0.05, **p<0.01, and ***p<0.001.

Next, we investigated the effect of ferristatin II-mediated TFRC inhibition on the expression of thermogenic markers. We found that ferristatin II prevented the db-cAMP-stimulated upregulation of UCP1 and PGC1a at both mRNA and protein levels in human SubQ and DeepC- derived adipocytes (**Fig. 5A, B**). In unstimulated condition, the inhibitor also reduced the protein expression of PGC1a in the DeepC-derived adipocytes with high thermogenic competency (**Fig. 5B**), likely because of impaired mitochondrial biogenesis resulting from iron deficiency. The adrenergic-driven upregulation of other thermogenic marker genes including *DIO2*, *CITED1*, and *PM20D1* was also hampered by ferristatin II (**Fig. 5C**). These findings suggest that iron also plays a role in heat generation by regulating gene expression of thermogenic markers.

**Figure 5.**
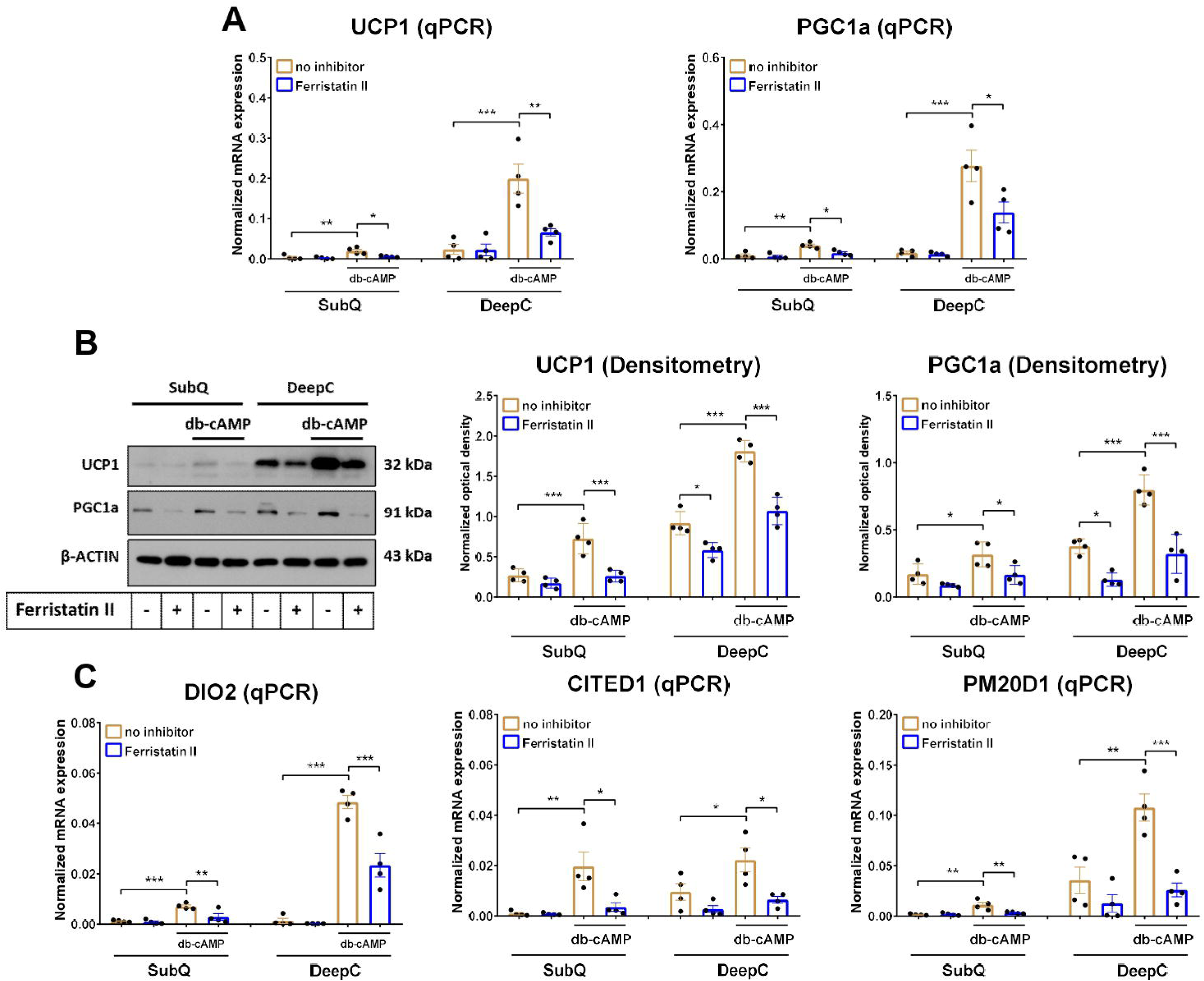
Inhibition of transferrin receptor (TFRC) by ferristatin II decreased the expression of thermogenic markers in human *ex vivo* differentiated subcutaneous (SubQ) and deep cervical (DeepC)-derived adipocytes. Adipocytes were differentiated and treated as in Figure 4. (A-B) mRNA (A) and protein (B) expression of UCP1 and PGC1a detected by RT- qPCR and immunoblotting, respectively. The original uncropped images of the full-length blots are displayed in **Supplementary Fig. 6.** (C) mRNA expression of *DIO2*, *CITED1*, and *PM20D1* detected by RT-qPCR. n=4, statistical analysis was performed by one-way ANOVA followed by Tukey’s *post hoc* test, *p<0.05, **p<0.01, and ***p<0.001.

### TFRC knock-down by siRNA impaired mitochondrial respiration and the expression of mitochondrial complex subunits and thermogenic markers during thermogenic activation of DeepC-derived adipocytes

To specifically examine the effects of TFRC, finally we used siRNA targeting multiple sequences to knock-down TFRC expression in the thermogenically prone DeepC-derived adipocytes. First, we checked whether the knock-down was successful and found that the mRNA and protein expression of TFRC was significantly reduced upon siRNA-mediated knock-down (**Fig. 6A, B**). In parallel, the db-cAMP-driven elevation of intracellular iron content (**Fig. 6C**), stimulated maximal, and proton leak respiration (**Fig. 6D**) was also prevented as a result of TFRC silencing. We also observed that the protein expression of mitochondrial complex subunits I, II, and IV was decreased in response to TFRC knock-down during adrenergic activation (**Fig. 6E**). Next, we investigated the expression of thermogenic genes upon TFRC silencing and found that the db- cAMP-stimulated upregulation of UCP1 and PGC1a was hampered by TFRC knock-down (**Fig. 7A, B**). In addition, the mRNA expression of *DIO2* and *CITED1* thermogenic markers was also decreased as a result of TFRC silencing during activation (**Fig. 7C**). These findings demonstrate that TFRC-mediated iron uptake is essential to support mitochondrial respiration, the expression of iron-containing respiratory chain proteins, and thermogenic genes expression during adrenergic stimulation in human thermogenic adipocytes.

**Figure 6.**
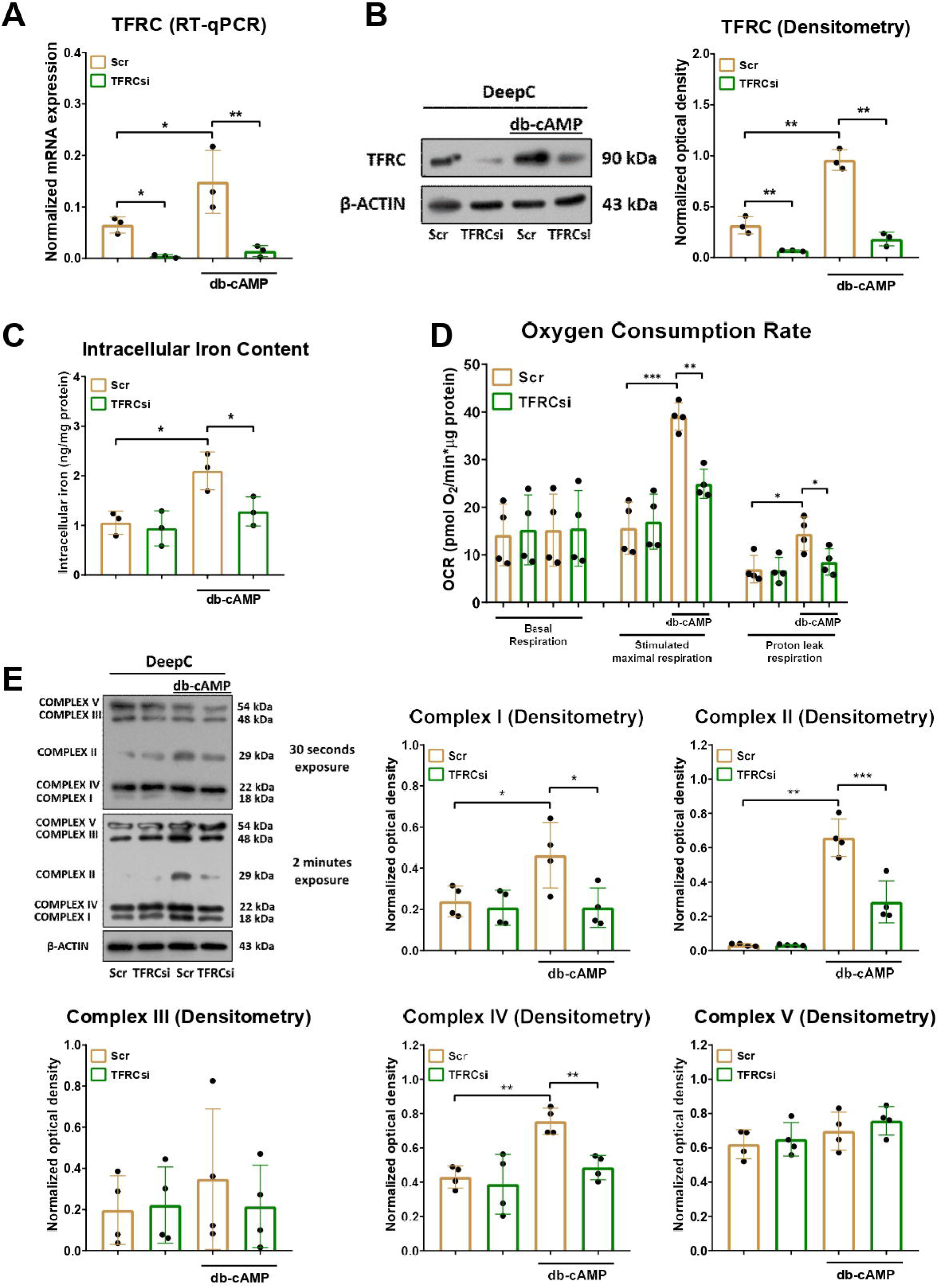
The effect of transferrin receptor (TFRC) knock-down on intracellular iron level, oxygen consumption, and protein expression of mitochondrial complex subunits in human *ex vivo* differentiated deep cervical (DeepC)-derived adipocytes. (A-B) mRNA (A) and protein expression (B) of TFRC upon silencing by siRNA detected by RT-qPCR and immunoblotting, n=3. (C) Intracellular iron level in TFRC-silenced DeepC-derived adipocytes during adrenergic stimulation for 10 h, n=3. (D) Basal, db-cAMP stimulated maximal, and proton leak oxygen consumption rate (OCR) in TFRC-silenced adipocytes were quantified by Seahorse extracellular flux analysis, n=4. (E) Protein expression of mitochondrial complex subunits detected by immunoblotting, n=4. The original uncropped images of the full-length blots are displayed in **Supplementary Fig. 7**. Statistical analysis was performed by one-way ANOVA followed by Tukey’s *post hoc* test, *p<0.05, *p<0.01, and ***p<0.001.

**Figure 7.**
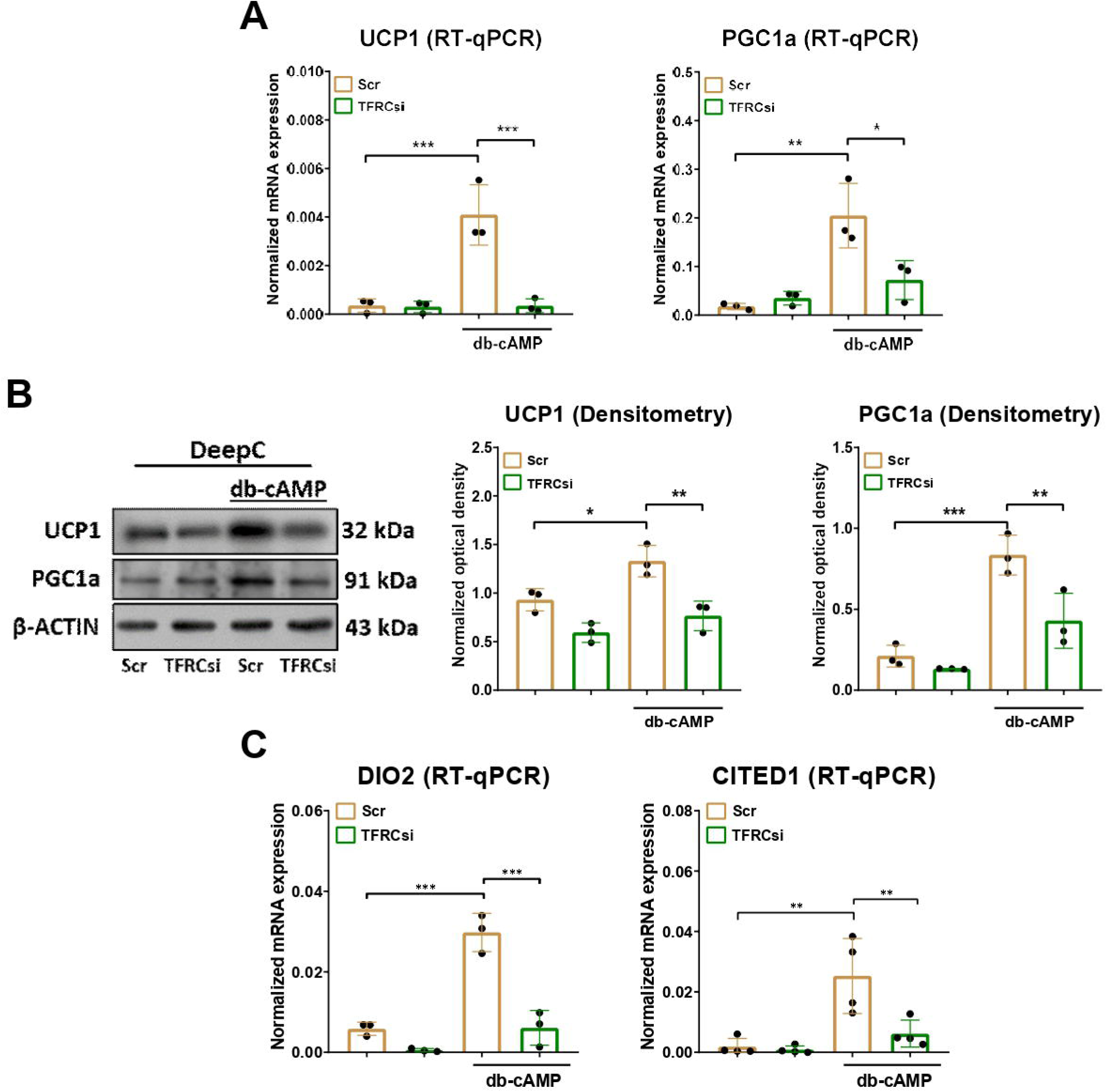
The effect of transferrin receptor (TFRC) knock-down on the expression of thermogenic markers in human *ex vivo* differentiated deep cervical (DeepC)-derived adipocytes. (A-B) mRNA (A) and protein (B) expression of UCP1 and PGC1a detected by RT-qPCR and immunoblotting, respectively. The original uncropped images of the full-length blots are displayed in **Supplementary Fig. 8.** (C) mRNA expression of *DIO2* and *CITED1* detected by RT-qPCR. n=4, statistical analysis was performed by one-way ANOVA followed by Tukey’s *post hoc* test, *p<0.05, **p<0.01, and ***p<0.001.

### Ferroportin inhibition did not increase further mitochondrial respiration and the expression of mitochondrial complex subunits and thermogenic markers in human cervical-derived adipocytes

Previous RNA-seq data was validated **(Fig. 1G)** showing that the expression of *SLC40A1* that encodes ferroportin was significantly decreased by db-cAMP at both mRNA and protein levels in SubQ and DeepC-derived adipocytes (**Supplementary Fig. 1A, B**). Ferroportin inhibitor, VIT- 2763, did not alter the expression of ferroportin at either mRNA or protein levels (**Supplementary Fig. 1A, B**). In parallel, the intracellular iron content (**Supplementary Fig. 1C**), oxygen consumption (**Supplementary Fig. 1D, E**), and mitochondrial complex subunits I-V expression (**Supplementary Fig. 1F**) were not affected significantly by VIT-2763 in neither control nor adrenergic stimulated conditions. We further assessed the effect of ferroportin inhibition on the expression of thermogenic markers and observed significant alterations neither in UCP1 and PGC1a mRNA or protein levels (**Supplementary Fig. 2A, B**) nor in the expression of other thermogenesis-related genes such as *DIO2*, *CITED1*, and *PM20D1* (**Supplementary Fig. 2C**). Collectively, these results suggest that ferroportin inhibition results in negligible influence on mitochondrial function or regulation of thermogenic gene expression in human cervical-derived adipocytes.

### Human thermogenic adipocytes expressed and secreted TF and MELTF

Having observed the importance of TFRC-mediated iron influx in the efficient thermogenic response of human adipocytes, we investigated its potential interacting partners including TF and MELTF. First, we utilized publicly available snRNA-seq database [Sun *et al*., 2020, *Nature*] and found that the expression of *TF* was enriched in adipocytes cluster within human BAT (**Fig. 8A**), while the expression of *MELTF* was detected in dendritic cells and preadipocytes (**Fig. 8B**) in the absence of thermogenesis activation. Consistently, analysis of isolated unstimulated human brown adipocytes from eight independent donors revealed that only the expression of *TF* was enriched (**Fig. 8C**) while *MELTF* was undetectable (**Fig. 8D**). These findings are in association with our bulk RNA-seq data in which we found that *TF* was constitutively expressed at a high level **(Fig. 1 E)** while the expression of *MELTF* was strongly induced upon adrenergic stimulation **(Fig. 1I)** in human adipocytes. Finally, we confirmed at the protein level that adipocytes secrete TF independently of stimulation (**Fig. 8E**), while the secretion of MELTF was elevated by adrenergic activation (**Fig. 8F**). Of note, MELTF secretion remained 20-50 times lower as compared to TF release even in db-cAMP stimulated condition suggesting that MELTF can support an auxiliary mechanism of iron uptake during active thermogenesis coupled with high iron demand.

**Figure 8.**
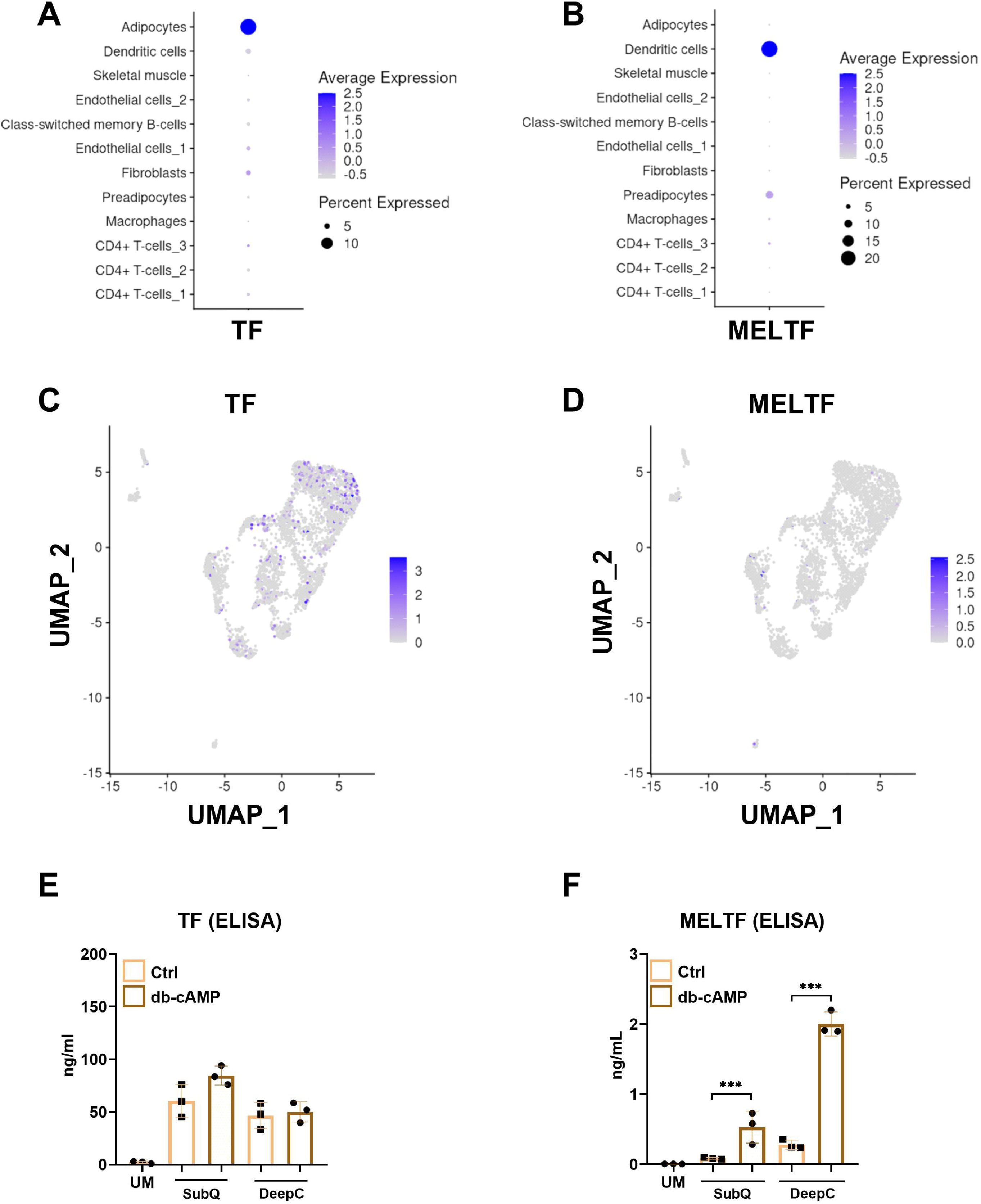
Transferrin (TF) and melanotransferrin (MELTF) were expressed and secreted by human adipocytes. (A, B) Dot plot displaying the expression of *TF* (A) and *MELTF* (B) in single-nuclei RNA-sequencing (snRNA-seq) data for human brown adipose tissue. (C, D) Embedding plots showing the expression of *TF* (C) and *MELTF* (D) by snRNA-seq from isolated brown adipocytes from 8 donors. Data displayed in panel A-D was retrieved from *batnetwork.org* [Sun *et al*., 2020, *Nature*]. (E, F) Protein secretion of TF (E) and MELTF (F) by control (ctrl) and dibutyryl (db)-cAMP-activated *ex vivo* differentiated subcutaneous (SubQ) and deep cervical (DeepC)-derived adipocytes detected by ELISA, n=3. Statistical analysis was performed by one-way ANOVA followed by Tukey’s *post hoc* test, ***p<0.001. UM: unconditioned medium.

### MELTF is predicted to form a complex with TF based on *in silico* analysis

The human TFRC is a homodimeric transmembrane protein which consists of intracellular, transmembrane, and extracellular domains [Testi *et al*., 2019] (**Fig. 9A**). The extracellular domain is responsible for binding TF; the receptor-ligand interaction occurs outside the cell leading to vesicular traffic into the cell. TF is a globular protein containing an N- and a C-terminal domain (N- and C-lobes, respectively). Both lobes can be divided into two subdomains: N1 and N2, as well as C1 and C2 [Silva *et al*., 2021]. Similarly to TF, MELTF also contains N- and C-lobes (and the respective N1, N2, C1, and C2 subdomains), however, the spatial arrangement of these domains is different that of the TF [Hayashi *et al*., 2021]. The N-lobes of TF and MELTF are highly analogous while the C-lobes exhibit relatively low overall structural similarity (**Fig. 9B**). However, it is important to note that despite the low similarity of the C-lobe domains, the C1 and C2 subdomains of TF and MELTF are highly similar (**Supplementary Fig. 9**). The arrangement of the domains and the subdomains might change due to the conformational flexibility provided by the interdomain hinge regions [Silva *et al*., 2021].

**Figure 9.**
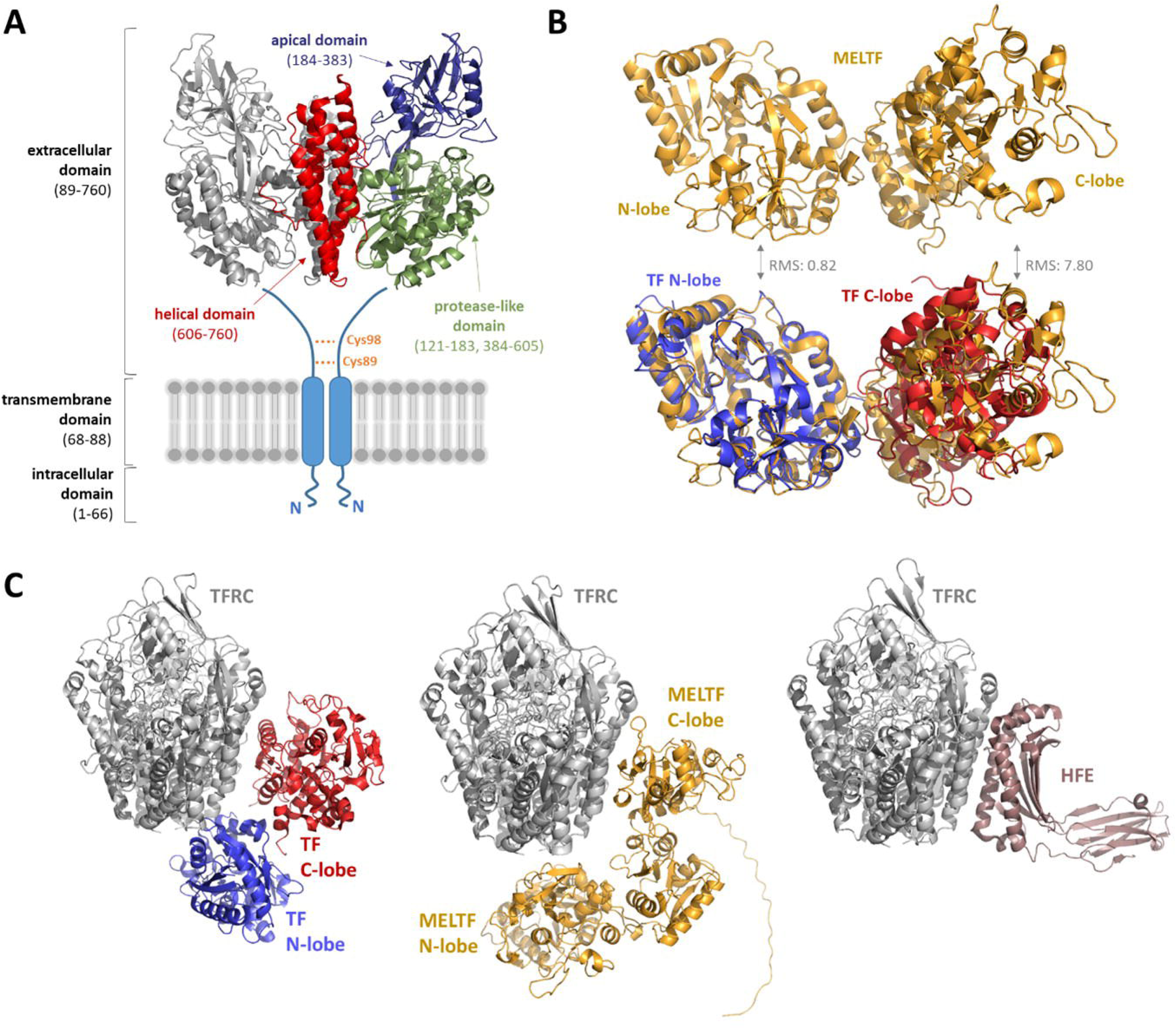
Structure of human transferrin receptor (TFRC) and its interactions with transferrin (TF), melanotransferrin (MELTF), and homeostatic iron regulator (HFE). (**A**) Overall structure of human TFRC. The domain organization and boundaries are shown based on UniProt database (UniProt ID: P02786) and Testi *et al*. [Testi *et al*., 2019]. The structure of the extracellular domain (122-756 residues) is shown by ribbon representation based on an electron microscopy structure (PDB ID: 1SUV) [Cheng *et al*., 2004]. The helical, apical, and protease-like domains are shown by different colors, the cysteine residues forming disulfide bonds are shown by yellow. (**B**) Structural alignment of TF and MELTF. TF and MELTF are shown by ribbon representation, based on experimentally determined structures. The complex of TFRC and TF is represented based on an electron microscopy structure (PDB ID: 1SUV) [Cheng *et al*., 2004]. The N- and C-lobes of TF have blue and red colors, respectively. The structure of MELTF complex is shown based on a crystal structure (PDB ID: 6XR0) [Hayashi *et al*., 2021]. (**C**) Binding of TF, MELTF, and HFE to TFRC. TFRC is shown in side-view and is colored by green, the interaction partners have different colors. The complex of TFRC and TF is represented based on an electron microscopy structure (PDB ID: 1SUV) [Cheng *et al*., 2004]. The N- and C-lobes of TF have blue and red colors, respectively. The complex of TFRC and MELTF is shown based on a proposed model that was prepared in this work by using AlphaFold. The binding of HFE to TFRC is shown based on a crystal structure (PDB ID: 1DE4) [Bennett *et al*., 2000]. RMS: Root Mean Square deviation.

The previously observed strong induction of MELTF secretion during thermogenesis activation prompted us to estimate the possible binding mode of MELTF to TFRC *in silico*. Model complexes were prepared (**Supplementary Fig. 10**) and the putative binding mode of MELTF was compared to that of TF and HFE proteins (**Fig. 9C**). Because of the structural similarity of the N-lobes, we assumed that binding of MELTF to TFRC might resemble that of TF, at least in the context of the N-lobe of MELTF. The predicted binding mode of MELTF was seemingly similar to that of the TF, the area of the interaction surfaces and the number of intermolecular contacts were also comparable (**Supplementary Table 3**). The proposed model showed that C-lobe of MELTF forms most of the polar interactions (H-bonds and salt bridges) with the helical domain of TFRC. Similar to this, TF and HFE also interact with the helical domain of TFRC (**Fig. 9C**). Despite the high structural similarity of the N-lobes of TF and MELTF **(Fig. 9B)**, the binding modes of these domains are not identical (**Fig. 9C**). The C-lobes exhibit considerable differences **(Fig. 9B)** and their binding mode is also different (**Fig. 9C**). Although the overall number of the interactions are comparable, the pattern of interactions between TFRC and MELTF is considerably different from that of the TF, only few TFRC residues contribute to binding both MELTF and TF via polar interactions (D126 of the protease-like domain as well as R623, R629, R646, E664, and E759 residues of the helical domain).

The comparison of the binding modes implies that MELTF, TF, and HFE also use the same surfaces and interact mainly with the helical domain of TFRC, but the pattern of interactions and the spatial positions of the subdomains are different. We predict that binding of MELTF to TFRC is not identical with that of the TF.

### TF and MELTF expression in human WAT positively correlated with metabolic health

As a final step, to explore the potential role of extracellular iron transport proteins in adipose tissue function, we examined the expression of TF and MELTF in human WAT and their association with metabolic health. When data from the open access adiposetissue.org database [Zhong *et al*., 2025, *Cell Metab.*] was retrieved, we found that the expression of *MELTF* in WAT was higher in non-obese individuals as compared to patients with obesity and it was significantly increased by weight loss [Salcedo-Tacuma *et al*., 2022, *Sci. Data*; Arner *et al*., 2018, *Cell Metab.*; Van Bussel *et al*., 2017, *Int. J. Obes.*; Petrus *et al*., 2018, *Cell Rep.*] (**Fig. 10A**). In addition, the mRNA expression of *MELTF* was elevated after 8 weeks of diet resulting in weight loss [Imbert *et* al. 2022, *J. Clin. Endocrinol. Metab.*; Armenise *et al*., 2017, *Am. J. Clin. Nutr.*] (**Fig. 10B**). We also found that the expression of *TF* in human abdominal SubQ WAT was inversely correlated with body mass index (BMI), waist and hip circumference, homeostatic model assessment of insulin resistance (HOMA-IR), circulating glucose and insulin, circulating low-density lipoprotein (LDL), total cholesterol, hemoglobin A1c (HbA1c), and fat cell volume, while positively correlated with blood plasma high-density lipoprotein levels [Zhong *et al*., 2025, *Cell Metab.*] (**Fig. 10C**). These findings suggest that the expression of TF and MELTF in human WAT is linked to improved metabolic health and corroborates the importance of iron transport in adipose tissue as a contributor to metabolic homeostasis.

**Figure 10.**
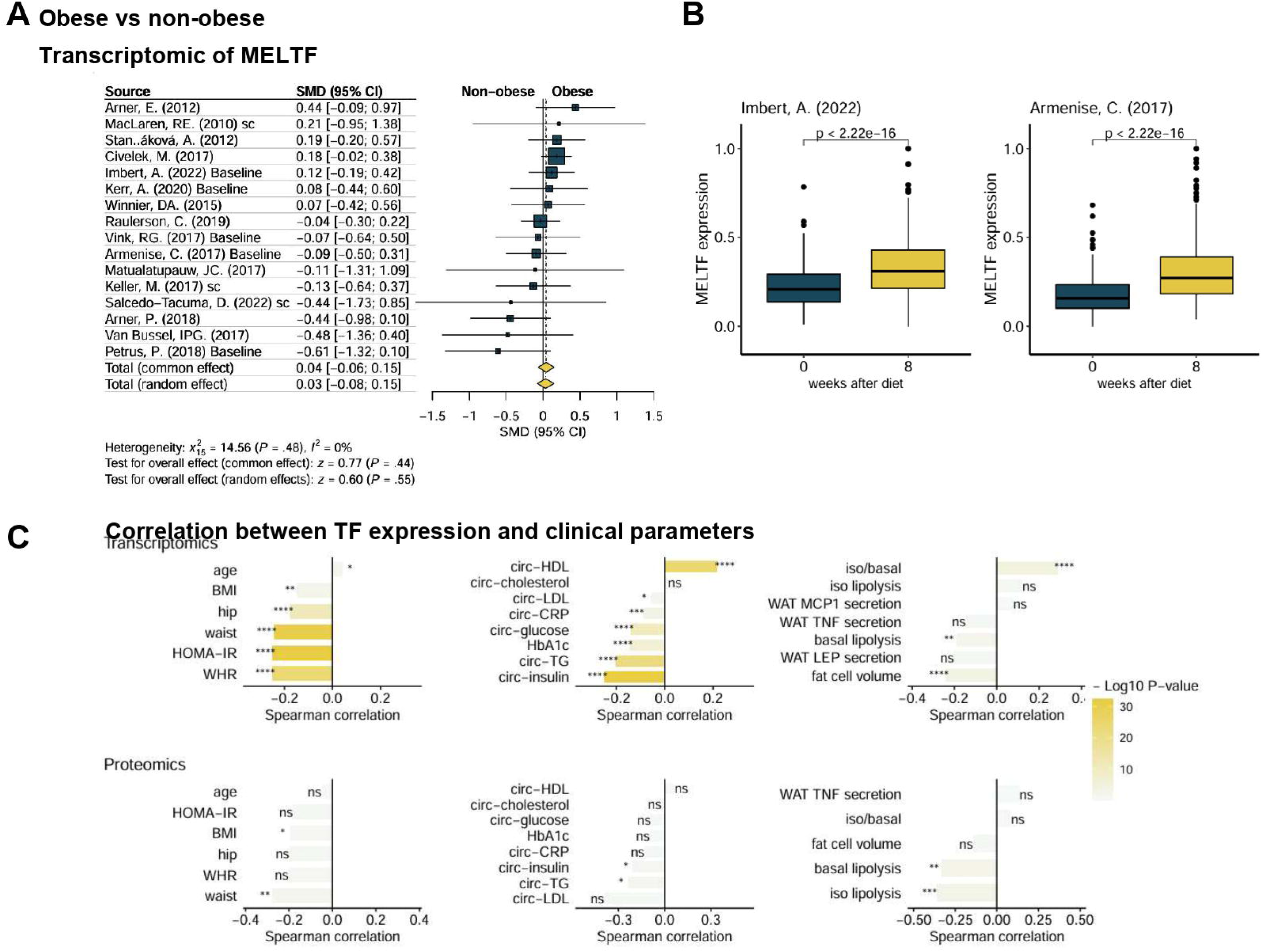
Correlation of melanotransferrin (MELTF) and transferrin (TF) expression in abdominal subcutaneous white adipose tissue (WAT) with clinical parameters. Data was retrieved from *adiposetissue.org* [Zhong *et al*., 2025]. (A) Meta-analysis forest plot comparing the *MELTF* mRNA levels in people living with or without obesity based on the respective BMI cutoffs (<30 kg/m^2^ vs. ≥30 kg/m^2^). SMD: Standardized mean differences [Data referenced from Arner *et al.,* 2012, *Diabetes*; Kerr *et al.,* 2020, *J. Internal Med.*; Petrus *et al.,* 2018, *Cell Rep.*; Arner *et al.,* 2018, *Cell Metab.;* Armenise *et al.,* 2017, *Am. J. Clin. Nutr.*; Salcedo-Tacuma *et al.,* 2022, *Scientific Data;* Civelek *et al.,* 2017, *Am. J Hum. Genet.*; Raulerson *et al.,* 2019, *Am. J Hum. Genet.;* Imbert *et al.,* 2022, *J. Clin. Endocrinol. Metab.*; Stančáková *et al.,* 2012, *Diabetes;* Winnier *et al.,* 2015, *Plos One;* Vink *et al.,* 2017, *Int. J Obes.;* MacLaren *et al.,* 2010, *BMC Med. Genom.*; Van Bussel *et al.,* 2017, *Int. J Obes;* Keller *et al.,* 2016, *Mol. Metab.*; and Matulalupauw *et al.,* 2017, *Int. J Obes*]. (B) Boxplots for 2 diet-induced weight loss cohorts indicate that *MELTF* mRNA expression in WAT was increased after the interventions (Data referenced from [*Cell Rep.*; Imbert *et al*., 2022, *J. Clin. Endocrinol.*, Armenise *et al*., 2017, *Am. J. Clin. Nutr*] (*adiposetissue.org*)). (C) The transcriptome (upper panels) and proteome (bottom panels) analyses were categorized into three groups: anthropometric (left panels), focusing on TF gene and protein expression in correlation with measurements of body distribution parameters (e.g., BMI); circulating diagnostic markers (middle panels), examining molecules present in the bloodstream (e.g., hormones); and tissue-specific responses (right panels), highlighting genes and proteins predominantly expressed in abdominal WAT. This classification helps to differentiate systemic metabolic regulation from localized tissue functions and their associations with body composition. Data in upper and bottom right panels were obtained from abdominal WAT-derived adipocytes differentiated *ex vivo*. BMI: body mass index; HOMA-IR: Homeostatic Model Assessment for Insulin Resistance; WHR: waist-hip ratio; HDL: high-density lipoprotein; HbA1c: hemoglobin A1c; LDL: low-density lipoprotein; TG: triglyceride; TNF: tumor necrosis factor; LEP: leptin; CRP: C-reactive protein; iso: isoproterenol.

## Discussion

Thermogenically active adipocytes require high amounts of nutrients including iron, in particular to support mitochondrial biogenesis and energy metabolism [Kim *et al*., 2022, *Clin. Nutr. Res.*]. The regulation of iron homeostasis has been recognized as a critical determinant for the thermogenic competency of murine adipocytes. Previous studies showed that DFO-mediated iron chelation impaired the browning program by modulating the expression of *Adipoq*, *Pparg*, *Ppargc1a*, and *Ucp1* in mouse 3T3-L1 cell line [Moreno-Navarette *et al*., 2014, *Diabetologia*] and in immortalized murine beige adipocytes [Qiu *et al*., 2020, *Front. Cell Dev. Biol.*]. Yook *et al*., (2021, *JBC*) also reported that the iron chelator treatment during differentiation significantly reduced the expression of Ucp1, Prdm16, CytC, Pgc1a, and mitochondrial complex subunits I, II, and IV in HIB1B-derived mouse brown adipocytes and C3H/10T1/2-derived murine beige adipocytes. Another study found that dietary iron deficiency disturbed iron homeostasis in mouse inguinal WAT (iWAT) and impaired adaptive thermogenesis, therefore, the mice became more prone to diet-induced weight gain [Yook *et al*., 2021, *J. Nutr.*].

Wang *et al*. (2017, *Biochem. Biophys. Res. Comm.*) described that iron content of mouse BAT was significantly increased upon acute cold challenge. However, the mechanisms by which the SNS-driven thermogenesis activation mediates iron accumulation in brown adipocytes, especially in human primary cell models, are still unrevealed. In our experiments, we validated that thermogenic adipocytes have elevated iron demands which are closely linked to mitochondrial biogenesis. During adrenergic stimulation by db-cAMP, human cervical-derived adipocytes significantly increased their intracellular iron levels along with the expression of TFRC and MELTF, while downregulated the expression of the iron exporter ferroportin, indicating that an expanded iron pool is required for supporting heat generation. We found that *ex vivo* iron depletion by DFO during adrenergic stimulation led to the decreased expression of thermogenic genes and mitochondrial complex subunits I, II, and IV, and oxygen consumption. Iron and heme participate in mitochondrial energy metabolism, as a component of the iron-sulfur clusters and cytochromes, including tricarboxylic acid cycle, oxidative phosphorylation, and fatty acid oxidation. Iron depletion, along with the reduced iron-sulfur cluster synthesis, results in impaired function of electron transport chain and thermogenesis, which then lead to decreased energy expenditure [Read *et al*., 2021, *Redox Biol.*; Belot *et al*., 2024, *Liver Int.*]. In addition to mitochondrial biogenesis, iron was also found to be essential for rewriting epigenetic marks during adipocyte differentiation by interacting with histone demethylase jumonji domain containing 1A (JMJD1A) and the DNA demethylase ten-eleven translocation 2 (TET2) [Suzuki *et al*., 2023] suggesting that it also plays important role in the regulation of gene expression. Future experiments are required to understand how the iron content of adipocytes can influence chromatin structure and the transcription of key genes mediating critical metabolic events along with thermogenesis when the cells are activated by an adrenergic cue.

TFRC is a cell surface receptor necessary for cellular iron uptake by the process of receptor-mediated endocytosis [Kawabata, 2019, *Free Radic. Biol. Med.*]. In accordance with our findings, a previous study also reported that the expression of Tfrc was induced by cold, β3- adrenergic agonist, and rosiglitazone in mouse iWAT and BAT. In contrast, high fat diet (HFD) and aging significantly decreased the expression of Tfrc in murine iWAT and BAT [Qiu *et al*., 2020, *Front. Cell. Dev. Biol.*]. Other studies in mice showed that the inhibition of Tfrc-mediated iron influx impaired mitochondrial function and beige adipocyte differentiation [Qiu *et al*., 2020, *Front. Cell. Dev. Biol.*; Li *et al*., 2020, *Adv. Sci.*; Yook *et al*., 2021, *JBC*]. In mice, Tfrc deficiency led to HFD-induced dyslipidemia, insulin resistance, and inflammation [Li *et al*., 2020, *Adv. Sci.*]. Partial reduction of Tfrc expression (around 20%) downregulated the expression of Ucp1 protein by 50% [Yook *et al*., 2021, *JBC*]. Our data showed that both pharmacological inhibition and siRNA-mediated knock-down of TFRC abrogated the db-cAMP-stimulated upregulation of thermogenic markers including UCP1 and PGC1a, as well as the proton leak respiration, which is associated with UCP1-dependent heat generation, and the expression of mitochondrial complex subunits I, II, and IV. Yook *et al*., (2021, *JBC*) observed that Tfrc silencing during HIB1B-derived brown adipocyte differentiation led to reduced *Ftl* (that encodes ferritin light chain) expression, however, our data showed that human cervical-derived adipocytes expressed negligible levels of *FTL* and *FTH* (that encodes ferritin heavy chain) and their expression was not affected by db-cAMP (Arianti *et al*., 2024, *Sci. Rep.*). Li *et al*. (2020, *Adv. Sci.*) proposed that *Tfrc* gene may be transcriptionally regulated by hypoxia-inducible factor 1 alpha (Hif1α) in mouse iWAT but not in BAT as they found that Hif1α was enriched at the *Tfrc* promoter during beigeing. Our data extended the current knowledge based on experiments carried out in mouse models by supporting that TFRC-mediated iron uptake is rapidly induced and it is a critical determinant for efficient thermogenic response in human BAT as a result of an SNS stimulus.

Ferroportin, encoded by *SLC40A1*, transports iron from the inside to the outside of the cell, playing a key role in maintaining systemic iron homeostasis [Mitchell *et al*., 2014, *Am J Physiol Cell Physiol.*]. A previous study reported that the expression of *SLC40A1* was significantly lower in thermogenic as compared to non-thermogenic adipocytes derived from human abdominal SubQ WAT [Min *et al*., 2019, *PNAS*]. Contrarily, adipose tissue expansion by overfeeding significantly elevated the expression of *SLC40A1* in abdominal SubQ WAT of healthy men [Segrestin *et al*., 2018, *JECM*]. In parallel, our data showed that adrenergic stimulation significantly decreased the expression of ferroportin. However, the pharmacological inhibition of ferroportin affected neither the expression of thermogenic markers nor oxygen consumption significantly. In contrast to our findings, β3-adrenergic receptor activation by CL-316243 increased the protein expression of ferroportin in mouse iWAT and this elevation was diminished when the stimulus was withdrawn [Yook *et al*., 2021, *PNAS*]. Another study showed that the lack of ferroportin in murine adipocytes resulted in iron accumulation and insulin resistance [Gabrielsen *et al*., 2012, *J. Clin. Invest.*]. On the other hand, Britton *et al*. (2018, *Cell Mol Gastroenterol Hepatol.)* found that adipocyte-specific ferroportin deletion did not lead to increased adipocyte iron stores and dysregulation of adiponectin, leptin, and resistin expression. The observed discrepancy between the current and previously published results may reflect the differences in species (humans vs mice), adipose tissue depots (cervical depots vs iWAT), and the complexity of iron metabolism regulation. Ferroportin is regulated by hepcidin, which binds to the iron exporter and promotes its internalization and degradation [Qiao *et al*., 2012, *Cell Metab.*]. Hepcidin, encoded by *HAMP*, is primarily secreted by hepatocytes. Extrahepatic production of hepcidin has been reported in kidney, heart, and adipose tissue, however, our RNA-sequencing data showed that the expression of *HAMP* was negligible in human cervical-derived adipocytes [Arianti *et al*., 2024, *Sci. Rep.*].

We also found that TF was secreted by *ex vivo* differentiated human primary adipocytes, however, db-cAMP-driven stimulation increased further neither its expression nor its release. SnRNA-seq data retrieved from Sun *et al*. (2020, *Nature*) showed that the expression of *TF* is abundant in the adipocyte cell clusters within human BAT originated from the DeepC depot even in the absence of adrenergic activation indicating that the released TF may play an active role in iron homeostasis and metabolic regulation within thermogenic adipocytes. The expression of TF in human abdominal SubQ WAT is inversely correlated with BMI, waist and hip circumference, HOMA-IR, adipocyte volume, and blood plasma glucose, insulin, LDL, total cholesterol, and HbA1c levels [Zhong *et al*., 2025, *Cell Metab.*].

MELTF is an iron-binding member of the TF superfamily which can be membrane-anchored or secreted to the serum [Hayashi *et al*., 2021, *Sci. Rep.*]. It is highly expressed in various malignant cells including melanoma, colorectal [Shin *et al*., 2014, *J. Proteom. Res.*], and lung cancers [Lei *et al*., 2020, *Cell Death and Disease*]. Hitherto, the function of MELTF in iron transport into adipocytes remained unclear. First, we found that the MELTF expression and secretion by adipocytes was strongly elevated by adrenergic stimulation of thermogenesis. By utilizing AlphaFold2 and HADDOCK, the predicted interaction surface for TFRC-MELTF was similar to that of TFRC-TF and TFRC-HFE, nevertheless, the pattern of receptor-ligand interactions, *i.e.* the mode of binding to TFRC is different. The predicted model complexes imply ability of MELTF for binding to TFRC and our computation analyses are in agreement with a previous assumption according to which the MELTF potentially has an ability for binding to TFRC, due to its high degree of homology with TF [Sekyere and Richardson, 2000, *FEBS Lett*]. These results suggest MELTF’s possible role in mediating iron influx (**Fig. 9**) to support adipocyte thermogenesis when the cells are activated by an adrenergic stimulus. When data from the open access adiposetissue.org database [Zhong *et al*., 2025, *Cell Metab.*] was retrieved, we found that the expression of *MELTF* in WAT was higher in non-obese individuals as compared to patients with obesity and it is significantly increased by weight loss [Salcedo-Tacuma *et al*., 2022, *Sci. Data*; Arner *et al*., 2018, *Cell Metab.*; Van Bussel *et al*., 2017, *Int. J. Obes.*; Petrus *et al*., 2018, *Cell Rep.*]. In addition, the mRNA expression of *MELTF* is elevated after 8 weeks of diet [Imbert *et* al. 2022, *J. Clin. Endocrinol. Metab.*; Armenise *et al*., 2017, *Am. J. Clin. Nutr.*]. The role of MELTF in iron uptake remains to be confirmed experimentally in distinct physiological or pathological situations. Previous studies reported that MELTF mediates iron uptake in the brain [Rothenberger *et al*., 1996, *Brain Research*; Jefferies *et al*., 1996, *Brain Research*], but not in a human melanoma cell line [Richardson and Baker, 1991, *BBA-Mol. Cel. Res.*]. Although our *in silico* analysis predicted binding between TFRC and MELTF, functional experiments are required to confirm this interaction and to delineate the iron transporting role of MELTF in the function of adipocytes, especially when they are stimulated by the SNS.

The relationship between obesity and iron deficiency was first observed six decades ago [Wenzel *et al*., 1962, *Lancet*; Seltzer and Mayer, 1963, *Am J. Clin. Nutr.*]. A quantitative meta-analysis study involving 13.393 individuals with obesity and 26.621 lean subjects reported that individuals with obesity had lower serum iron levels and higher risk of iron deficiency as compared to lean ones [Zhao *et al*., 2015, *Obes. Rev.*]. A recent Mendelian randomization study supported a causal link between obesity and iron deficient anemia and strengthened the epidemiological evidence that iron metabolism is impaired in obesity [Wang *et al*., 2023, *Front. Public Health*]. Iron deficiency causes fatigue and in severe cases immunological, developmental, or neurocognitive defects [Pantopoulos, 2024, *Haematologica*]. Oral iron supplementation is commonly used as first-line therapy due to its convenience and low cost, but effective repletion requires relatively high doses (50–200 mg/day for 3–12 weeks) of which only ∼10–20% is absorbed and therefore can trigger side effects [Celis *et al*., 2023, *Cell Chem. Biol.*]. Careful assessment is required before recommending iron supplementation for the prevention or treatment of obesity-related metabolic disorders. In cases where obesity is accompanied by iron deficiency, supplementation may help to restore adipocyte function and thereby contribute to improved metabolic health at least partially by supporting thermogenic activation. Taken together, our findings provide supporting evidence that iron is a critical nutritional factor in stimulating adipocyte thermogenesis that leads to augmented energy expenditure to combat obesity.

## Author contributions

R. Al. — methodology, investigation, data curation, validation, visualization, writing – original draft; M. S — methodology, investigation, data curation; G. K. — investigation; F. R. M. — methodology, formal analysis, software; M.Á.D — methodology, formal analysis, software; F. G. — methodology, resources; L. F. — conceptualization, funding acquisition, supervision, writing – review and editing; J. A. M. — methodology, formal analysis, software; E. K. — conceptualization, project administration, funding acquisition, supervision, writing – review and editing; R.Ar. — conceptualization, formal analysis, data curation, validation, visualization, writing - original draft.

## Acknowledgements

We thank Dr. Éva Csősz for the exceptional help in reviewing the manuscript before its submission, Boglárka Ágnes Vinnai and Yousif Qais Al-Khafaji for cell culture and RT-qPCR assays, and Jennifer Nagy for technical assistance.

## References

Arianti, R., Vinnai, B. Á., Alrifai, R., Karadsheh, G., Al-Khafaji, Y. Q., Póliska, S., Győry, F., Fésüs, L., & Kristóf, E. (2024). Upregulation of inhibitor of DNA binding 1 and 3 is important for efficient thermogenic response in human adipocytes. Scientific reports, 14(1), 28272. 10.1038/s41598-024-79634-2

Arianti, R., Vinnai, B. Á., Győry, F., Guba, A., Csősz, É., Kristóf, E., & Fésüs, L. (2023). Availability of abundant thiamine determines efficiency of thermogenic activation in human neck area derived adipocytes. The Journal of nutritional biochemistry, 119, 109385. 10.1016/j.jnutbio.2023.109385

Armenise, C., Lefebvre, G., Carayol, J., Bonnel, S., Bolton, J., Di Cara, A., Gheldof, N., Descombes, P., Langin, D., Saris, W. H., Astrup, A., Hager, J., Viguerie, N., & Valsesia, A. (2017). Transcriptome profiling from adipose tissue during a low-calorie diet reveals predictors of weight and glycemic outcomes in obese, nondiabetic subjects. The American journal of clinical nutrition, 106(3), 736–746. 10.3945/ajcn.117.156216

Arner, E., Mejhert, N., Kulyté, A., Balwierz, P. J., Pachkov, M., Cormont, M., Lorente-Cebrián, S., Ehrlund, A., Laurencikiene, J., Hedén, P., Dahlman-Wright, K., Tanti, J. F., Hayashizaki, Y., Rydén, M., Dahlman, I., van Nimwegen, E., Daub, C. O., & Arner, P. (2012). Adipose tissue microRNAs as regulators of CCL2 production in human obesity. Diabetes, 61(8), 1986–1993. 10.2337/db11-1508

Arner, P., Andersson, D. P., Bäckdahl, J., Dahlman, I., & Rydén, M. (2018). Weight Gain and Impaired Glucose Metabolism in Women Are Predicted by Inefficient Subcutaneous Fat Cell Lipolysis. Cell metabolism, 28(1), 45–54.e3. 10.1016/j.cmet.2018.05.004

Belot, A., Puy, H., Hamza, I., & Bonkovsky, H. L. (2024). Update on heme biosynthesis, tissue-specific regulation, heme transport, relation to iron metabolism and cellular energy. Liver international : official journal of the International Association for the Study of the Liver, 44(9), 2235–2250. 10.1111/liv.15965

Bennett, M. J., Lebrón, J. A., & Bjorkman, P. J. (2000). Crystal structure of the hereditary haemochromatosis protein HFE complexed with transferrin receptor. Nature, 403(6765), 46–53. 10.1038/47417

Britton, L., Jaskowski, L. A., Bridle, K., Secondes, E., Wallace, D., Santrampurwala, N., Reiling, J., Miller, G., Mangiafico, S., Andrikopoulos, S., Subramaniam, V. N., & Crawford, D. (2018). Ferroportin Expression in Adipocytes Does Not Contribute to Iron Homeostasis or Metabolic Responses to a High Calorie Diet. Cellular and molecular gastroenterology and hepatology, 5(3), 319–331. 10.1016/j.jcmgh.2018.01.005

Byrne, S. L., Buckett, P. D., Kim, J., Luo, F., Sanford, J., Chen, J., Enns, C., & Wessling-Resnick, M. (2013). Ferristatin II promotes degradation of transferrin receptor-1 in vitro and in vivo. PloS one, 8(7), e70199. 10.1371/journal.pone.0070199

Cannon, B., & Nedergaard, J. (2004). Brown adipose tissue: function and physiological significance. Physiological reviews, 84(1), 277–359. 10.1152/physrev.00015.2003

Celis, A. I., Relman, D. A., & Huang, K. C. (2023). The impact of iron and heme availability on the healthy human gut microbiome in vivo and in vitro. Cell chemical biology, 30(1), 110– 126.e3. 10.1016/j.chembiol.2022.12.001

Cheng, Y., Zak, O., Aisen, P., Harrison, S. C., & Walz, T. (2004). Structure of the human transferrin receptor-transferrin complex. Cell, 116(4), 565–576. 10.1016/s0092-8674(04)00130-8

Civelek, M., Wu, Y., Pan, C., Raulerson, C. K., Ko, A., He, A., Tilford, C., Saleem, N. K., Stančáková, A., Scott, L. J., Fuchsberger, C., Stringham, H. M., Jackson, A. U., Narisu, N., Chines, P. S., Small, K. S., Kuusisto, J., Parks, B. W., Pajukanta, P., Kirchgessner, T., … Lusis, A. J. (2017). Genetic Regulation of Adipose Gene Expression and Cardio-Metabolic Traits. American journal of human genetics, 100(3), 428–443. 10.1016/j.ajhg.2017.01.027

Cypess, A. M., Lehman, S., Williams, G., Tal, I., Rodman, D., Goldfine, A. B., Kuo, F. C., Palmer, E. L., Tseng, Y. H., Doria, A., Kolodny, G. M., & Kahn, C. R. (2009). Identification and importance of brown adipose tissue in adult humans. The New England journal of medicine, 360(15), 1509–1517. 10.1056/NEJMoa0810780

Gabrielsen, J. S., Gao, Y., Simcox, J. A., Huang, J., Thorup, D., Jones, D., Cooksey, R. C., Gabrielsen, D., Adams, T. D., Hunt, S. C., Hopkins, P. N., Cefalu, W. T., & McClain, D. A. (2012). Adipocyte iron regulates adiponectin and insulin sensitivity. The Journal of clinical investigation, 122(10), 3529–3540. 10.1172/JCI44421

Galy, B., Conrad, M., & Muckenthaler, M. (2024). Mechanisms controlling cellular and systemic iron homeostasis. Nature reviews. Molecular cell biology, 25(2), 133–155. 10.1038/s41580-023-00648-1

Hayashi, K., Longenecker, K. L., Liu, Y. L., Faust, B., Prashar, A., Hampl, J., Stoll, V., & Vivona, S. (2021). Complex of human Melanotransferrin and SC57.32 Fab fragment reveals novel interdomain arrangement with ferric N-lobe and open C-lobe. Scientific reports, 11(1), 566. 10.1038/s41598-020-79090-8

Honorato, R. V., Trellet, M. E., Jiménez-García, B., Schaarschmidt, J. J., Giulini, M., Reys, V., Koukos, P. I., Rodrigues, J. P. G. L. M., Karaca, E., van Zundert, G. C. P., Roel-Touris, J., van Noort, C. W., Jandová, Z., Melquiond, A. S. J., & Bonvin, A. M. J. J. (2024). The HADDOCK2.4 web server for integrative modeling of biomolecular complexes. Nature protocols, 19(11), 3219– 3241. 10.1038/s41596-024-01011-0

Huang, Z., Gu, C., Zhang, Z., Arianti, R., Swaminathan, A., Tran, K., Battist, A., Kristóf, E., & Ruan, H. B. (2023). Supraclavicular brown adipocytes originate from Tbx1+ myoprogenitors. PLoS biology, 21(12), e3002413. 10.1371/journal.pbio.3002413

Ibrahim, T., Khandare, V., Mirkin, F. G., Tumtas, Y., Bubeck, D., & Bozkurt, T. O. (2023). AlphaFold2-multimer guided high-accuracy prediction of typical and atypical ATG8-binding motifs. PLoS biology, 21(2), e3001962. 10.1371/journal.pbio.3001962

Imbert, A., Vialaneix, N., Marquis, J., Vion, J., Charpagne, A., Metairon, S., Laurens, C., Moro, C., Boulet, N., Walter, O., Lefebvre, G., Hager, J., Langin, D., Saris, W. H. M., Astrup, A., Viguerie, N., & Valsesia, A. (2022). Network Analyses Reveal Negative Link Between Changes in Adipose Tissue GDF15 and BMI During Dietary-induced Weight Loss. The Journal of clinical endocrinology and metabolism, 107(1), e130–e142. 10.1210/clinem/dgab621

Jefferies, W. A., Food, M. R., Gabathuler, R., Rothenberger, S., Yamada, T., Yasuhara, O., & McGeer, P. L. (1996). Reactive microglia specifically associated with amyloid plaques in Alzheimer’s disease brain tissue express melanotransferrin. Brain research, 712(1), 122–126. 10.1016/0006-8993(95)01407-1

Jespersen, N. Z., Larsen, T. J., Peijs, L., Daugaard, S., Homøe, P., Loft, A., de Jong, J., Mathur, N., Cannon, B., Nedergaard, J., Pedersen, B. K., Møller, K., & Scheele, C. (2013). A classical brown adipose tissue mRNA signature partly overlaps with brite in the supraclavicular region of adult humans. Cell metabolism, 17(5), 798–805. 10.1016/j.cmet.2013.04.011

Kajimura, S., Spiegelman, B. M., & Seale, P. (2015). Brown and Beige Fat: Physiological Roles beyond Heat Generation. Cell metabolism, 22(4), 546–559. 10.1016/j.cmet.2015.09.007

Kawabata H. (2019). Transferrin and transferrin receptors update. Free radical biology & medicine, 133, 46–54. 10.1016/j.freeradbiomed.2018.06.037

Keller, M., Hopp, L., Liu, X., Wohland, T., Rohde, K., Cancello, R., Klös, M., Bacos, K., Kern, M., Eichelmann, F., Dietrich, A., Schön, M. R., Gärtner, D., Lohmann, T., Dreßler, M., Stumvoll, M., Kovacs, P., DiBlasio, A. M., Ling, C., Binder, H., … Böttcher, Y. (2016). Genome-wide DNA promoter methylation and transcriptome analysis in human adipose tissue unravels novel candidate genes for obesity. Molecular metabolism, 6(1), 86–100. 10.1016/j.molmet.2016.11.003

Kerr, A. G., Andersson, D. P., Rydén, M., Arner, P., & Dahlman, I. (2020). Long-term changes in adipose tissue gene expression following bariatric surgery. Journal of internal medicine, 288(2), 219–233. 10.1111/joim.13066

Kim, S. L., Shin, S., & Yang, S. J. (2022). Iron Homeostasis and Energy Metabolism in Obesity. Clinical nutrition research, 11(4), 316–330. 10.7762/cnr.2022.11.4.316

Kremer, D. M., Nelson, B. S., Lin, L., Yarosz, E. L., Halbrook, C. J., Kerk, S. A., Sajjakulnukit, P., Myers, A., Thurston, G., Hou, S. W., Carpenter, E. S., Andren, A. C., Nwosu, Z. C., Cusmano, N., Wisner, S., Mbah, N. E., Shan, M., Das, N. K., Magnuson, B., Little, A. C., … Lyssiotis, C. A. (2021). GOT1 inhibition promotes pancreatic cancer cell death by ferroptosis. Nature communications, 12(1), 4860. 10.1038/s41467-021-24859-2

Laskowski, R. A., Jabłońska, J., Pravda, L., Vařeková, R. S., & Thornton, J. M. (2018). PDBsum: Structural summaries of PDB entries. Protein science : a publication of the Protein Society, 27(1), 129–134. 10.1002/pro.3289

Lei, Y., Lu, Z., Huang, J., Zang, R., Che, Y., Mao, S., Fang, L., Liu, C., Wang, X., Zheng, S., Sun, N., & He, J. (2020). The membrane-bound and soluble form of melanotransferrin function independently in the diagnosis and targeted therapy of lung cancer. Cell death & disease, 11(10), 933. 10.1038/s41419-020-03124-2

Leitner, B. P., Huang, S., Brychta, R. J., Duckworth, C. J., Baskin, A. S., McGehee, S., Tal, I., Dieckmann, W., Gupta, G., Kolodny, G. M., Pacak, K., Herscovitch, P., Cypess, A. M., & Chen, K. Y. (2017). Mapping of human brown adipose tissue in lean and obese young men. Proceedings of the National Academy of Sciences of the United States of America, 114(32), 8649–8654. 10.1073/pnas.1705287114

Li, J., Pan, X., Pan, G., Song, Z., He, Y., Zhang, S., Ye, X., Yang, X., Xie, E., Wang, X., Mai, X., Yin, X., Tang, B., Shu, X., Chen, P., Dai, X., Tian, Y., Yao, L., Han, M., Xu, G., … Xie, L. (2020). Transferrin Receptor 1 Regulates Thermogenic Capacity and Cell Fate in Brown/Beige Adipocytes. *Advanced science (Weinheim, Baden-Wurttemberg*, Germany), 7(12), 1903366. 10.1002/advs.201903366

MacLaren, R. E., Cui, W., Lu, H., Simard, S., & Cianflone, K. (2010). Association of adipocyte genes with ASP expression: a microarray analysis of subcutaneous and omental adipose tissue in morbidly obese subjects. BMC medical genomics, 3, 3. 10.1186/1755-8794-3-3

Matualatupauw, J. C., Bohl, M., Gregersen, S., Hermansen, K., & Afman, L. A. (2017). Dietary medium-chain saturated fatty acids induce gene expression of energy metabolism-related pathways in adipose tissue of abdominally obese subjects. International journal of obesity (2005), *41*(9), 1348–1354. 10.1038/ijo.2017.120

Min, S. Y., Desai, A., Yang, Z., Sharma, A., DeSouza, T., Genga, R. M. J., Kucukural, A., Lifshitz, L. M., Nielsen, S., Scheele, C., Maehr, R., Garber, M., & Corvera, S. (2019). Diverse repertoire of human adipocyte subtypes develops from transcriptionally distinct mesenchymal progenitor cells. Proceedings of the National Academy of Sciences of the United States of America, 116(36), 17970–17979. 10.1073/pnas.1906512116

Mirdita, M., Schütze, K., Moriwaki, Y., Heo, L., Ovchinnikov, S., & Steinegger, M. (2022). ColabFold: making protein folding accessible to all. Nature methods, 19(6), 679–682. 10.1038/s41592-022-01488-1

Mitchell, C. J., Shawki, A., Ganz, T., Nemeth, E., & Mackenzie, B. (2014). Functional properties of human ferroportin, a cellular iron exporter reactive also with cobalt and zinc. American journal of physiology. Cell physiology, 306(5), C450–C459. 10.1152/ajpcell.00348.2013

Moreno-Navarrete, J. M., Ortega, F., Moreno, M., Ricart, W., & Fernández-Real, J. M. (2014). Fine-tuned iron availability is essential to achieve optimal adipocyte differentiation and mitochondrial biogenesis. Diabetologia, 57(9), 1957–1967. 10.1007/s00125-014-3298-5

Negi, S. S., Schein, C. H., Oezguen, N., Power, T. D., & Braun, W. (2007). InterProSurf: a web server for predicting interacting sites on protein surfaces. *Bioinformatics (Oxford*, England), 23(24), 3397–3399. 10.1093/bioinformatics/btm474

Pantopoulos K. (2024). Oral iron supplementation: new formulations, old questions. Haematologica, 109(9), 2790–2801. 10.3324/haematol.2024.284967

Petrus, P., Mejhert, N., Corrales, P., Lecoutre, S., Li, Q., Maldonado, E., Kulyté, A., Lopez, Y., Campbell, M., Acosta, J. R., Laurencikiene, J., Douagi, I., Gao, H., Martínez-Álvarez, C., Hedén, P., Spalding, K. L., Vidal-Puig, A., Medina-Gomez, G., Arner, P., & Rydén, M. (2018). Transforming Growth Factor-β3 Regulates Adipocyte Number in Subcutaneous White Adipose Tissue. Cell reports, 25(3), 551–560.e5. 10.1016/j.celrep.2018.09.069

Qiao, B., Sugianto, P., Fung, E., Del-Castillo-Rueda, A., Moran-Jimenez, M. J., Ganz, T., & Nemeth, E. (2012). Hepcidin-induced endocytosis of ferroportin is dependent on ferroportin ubiquitination. Cell metabolism, 15(6), 918–924. 10.1016/j.cmet.2012.03.018

Qiu, J., Zhang, Z., Wang, S., Chen, Y., Liu, C., Xu, S., Wang, D., Su, J., Ni, M., Yu, J., Cui, X., Ma, L., Hu, T., Hu, Y., Gu, X., Ma, X., Wang, J., & Xu, L. (2020). Transferrin Receptor Functionally Marks Thermogenic Adipocytes. Frontiers in cell and developmental biology, 8, 572459. 10.3389/fcell.2020.572459

Raulerson, C. K., Ko, A., Kidd, J. C., Currin, K. W., Brotman, S. M., Cannon, M. E., Wu, Y., Spracklen, C. N., Jackson, A. U., Stringham, H. M., Welch, R. P., Fuchsberger, C., Locke, A. E., Narisu, N., Lusis, A. J., Civelek, M., Furey, T. S., Kuusisto, J., Collins, F. S., Boehnke, M., … Mohlke, K. L. (2019). Adipose Tissue Gene Expression Associations Reveal Hundreds of Candidate Genes for Cardiometabolic Traits. American journal of human genetics, 105(4), 773– 787. 10.1016/j.ajhg.2019.09.001

Read, A. D., Bentley, R. E., Archer, S. L., & Dunham-Snary, K. J. (2021). Mitochondrial iron-sulfur clusters: Structure, function, and an emerging role in vascular biology. Redox biology, 47, 102164. 10.1016/j.redox.2021.102164

Richardson, D., & Baker, E. (1991). The uptake of inorganic iron complexes by human melanoma cells. Biochimica et biophysica acta, 1093(1), 20–28. 10.1016/0167-4889(91)90133-i

Rosen, E. D., & Spiegelman, B. M. (2014). What we talk about when we talk about fat. Cell, 156(1-2), 20–44. 10.1016/j.cell.2013.12.012

Rothenberger, S., Food, M. R., Gabathuler, R., Kennard, M. L., Yamada, T., Yasuhara, O., McGeer, P. L., & Jefferies, W. A. (1996). Coincident expression and distribution of melanotransferrin and transferrin receptor in human brain capillary endothelium. Brain research, 712(1), 117–121. 10.1016/0006-8993(96)88505-2

Sakers, A., De Siqueira, M. K., Seale, P., & Villanueva, C. J. (2022). Adipose-tissue plasticity in health and disease. Cell, 185(3), 419–446. 10.1016/j.cell.2021.12.016

Salcedo-Tacuma, D., Bonilla, L., Montes, M. C. G., Gonzalez, J. E. N., Gutierrez, S. M. S., Chirivi, M., & Contreras, G. A. (2022). Transcriptome dataset of omental and subcutaneous adipose tissues from gestational diabetes patients. Scientific data, 9(1), 344. 10.1038/s41597-022-01457-5

Segrestin, B., Moreno-Navarrete, J. M., Seyssel, K., Alligier, M., Meugnier, E., Nazare, J. A., Vidal, H., Fernandez-Real, J. M., & Laville, M. (2019). Adipose Tissue Expansion by Overfeeding Healthy Men Alters Iron Gene Expression. The Journal of clinical endocrinology and metabolism, 104(3), 688–696. 10.1210/jc.2018-01169

Sekyere, E., & Richardson, D. R. (2000). The membrane-bound transferrin homologue melanotransferrin: roles other than iron transport?. FEBS letters, 483(1), 11–16. 10.1016/s0014-5793(00)02079-2

Seltzer, C. C., & Mayer, J. (1963). SERUM IRON AND IRON-BINDING CAPACITY IN ADOLESCENTS. II. COMPARISON OF OBESE AND NONOBESE SUBJECTS. The American journal of clinical nutrition, 13, 354–361. 10.1093/ajcn/13.6.354

Shamsi, F., Wang, C. H., & Tseng, Y. H. (2021). The evolving view of thermogenic adipocytes - ontogeny, niche and function. Nature reviews. Endocrinology, 17(12), 726–744. 10.1038/s41574-021-00562-6

Shin, J., Kim, H. J., Kim, G., Song, M., Woo, S. J., Lee, S. T., Kim, H., & Lee, C. (2014). Discovery of melanotransferrin as a serological marker of colorectal cancer by secretome analysis and quantitative proteomics. Journal of proteome research, 13(11), 4919–4931. 10.1021/pr500790f

Silva, A. M. N., Moniz, T., de Castro, B., Rangel, M. (2021). Human transferrin: An inorganic biochemistry perspective. Coordination Chemistry Reviews, 449, 214186. 10.1016/j.ccr.2021.214186

Sun, W., Dong, H., Balaz, M., Slyper, M., Drokhlyansky, E., Colleluori, G., Giordano, A., Kovanicova, Z., Stefanicka, P., Balazova, L., Ding, L., Husted, A. S., Rudofsky, G., Ukropec, J., Cinti, S., Schwartz, T. W., Regev, A., & Wolfrum, C. (2020). snRNA-seq reveals a subpopulation of adipocytes that regulates thermogenesis. Nature, 587(7832), 98–102. 10.1038/s41586-020-2856-x

Suzuki, T., Komatsu, T., Shibata, H., Tanioka, A., Vargas, D., Kawabata-Iwakawa, R., Miura, F., Masuda, S., Hayashi, M., Tanimura-Inagaki, K., Morita, S., Kohmaru, J., Adachi, K., Tobo, M., Obinata, H., Hirayama, T., Kimura, H., Sakai, J., Nagasawa, H., Itabashi, H., … Inagaki, T. (2023). Crucial role of iron in epigenetic rewriting during adipocyte differentiation mediated by JMJD1A and TET2 activity. Nucleic acids research, 51(12), 6120–6142. 10.1093/nar/gkad342

Stancáková, A., Civelek, M., Saleem, N. K., Soininen, P., Kangas, A. J., Cederberg, H., Paananen, J., Pihlajamäki, J., Bonnycastle, L. L., Morken, M. A., Boehnke, M., Pajukanta, P., Lusis, A. J., Collins, F. S., Kuusisto, J., Ala-Korpela, M., & Laakso, M. (2012). Hyperglycemia and a common variant of GCKR are associated with the levels of eight amino acids in 9,369 Finnish men. Diabetes, 61(7), 1895–1902. 10.2337/db11-1378

Szatmári-Tóth, M., Shaw, A., Csomós, I., Mocsár, G., Fischer-Posovszky, P., Wabitsch, M., Balajthy, Z., Lányi, C., Győry, F., Kristóf, E., & Fésüs, L. (2020). Thermogenic Activation Downregulates High Mitophagy Rate in Human Masked and Mature Beige Adipocytes. International journal of molecular sciences, 21(18), 6640. 10.3390/ijms21186640

Szklarczyk, D., Kirsch, R., Koutrouli, M., Nastou, K., Mehryary, F., Hachilif, R., Gable, A. L., Fang, T., Doncheva, N. T., Pyysalo, S., Bork, P., Jensen, L. J., & von Mering, C. (2023). The STRING database in 2023: protein-protein association networks and functional enrichment analyses for any sequenced genome of interest. Nucleic acids research, 51(D1), D638–D646. 10.1093/nar/gkac1000

Testi, C., Boffi, A., & Montemiglio, L. C. (2019). Structural analysis of the transferrin receptor multifaceted ligand(s) interface. Biophysical chemistry, 254, 106242. 10.1016/j.bpc.2019.106242

Tóth, B. B., Arianti, R., Shaw, A., Vámos, A., Veréb, Z., Póliska, S., Győry, F., Bacso, Z., Fésüs, L., & Kristóf, E. (2020). FTO Intronic SNP Strongly Influences Human Neck Adipocyte Browning Determined by Tissue and PPARγ Specific Regulation: A Transcriptome Analysis. Cells, 9(4), 987. 10.3390/cells9040987

Townsend, K. L., & Tseng, Y. H. (2014). Brown fat fuel utilization and thermogenesis. Trends in endocrinology and metabolism: TEM, 25(4), 168–177. 10.1016/j.tem.2013.12.004

Van Bussel, I. P. G., Backx, E. M. P., De Groot, C. P. G. M., Tieland, M., Müller, M., & Afman, L. A. (2017). The impact of protein quantity during energy restriction on genome-wide gene expression in adipose tissue of obese humans. International journal of obesity (2005), *41*(7), 1114–1120. 10.1038/ijo.2017.76

van Marken Lichtenbelt, W. D., & Schrauwen, P. (2011). Implications of nonshivering thermogenesis for energy balance regulation in humans. American journal of physiology. Regulatory, integrative and comparative physiology, 301(2), R285–R296. 10.1152/ajpregu.00652.2010

Vink, R. G., Roumans, N. J., Fazelzadeh, P., Tareen, S. H., Boekschoten, M. V., van Baak, M. A., & Mariman, E. C. (2017). Adipose tissue gene expression is differentially regulated with different rates of weight loss in overweight and obese humans. International journal of obesity (2005), *41*(2), 309–316. 10.1038/ijo.2016.201

Virtanen, K. A., Lidell, M. E., Orava, J., Heglind, M., Westergren, R., Niemi, T., Taittonen, M., Laine, J., Savisto, N. J., Enerbäck, S., & Nuutila, P. (2009). Functional brown adipose tissue in healthy adults. The New England journal of medicine, 360(15), 1518–1525. 10.1056/NEJMoa0808949

Vinnai, B. Á., Arianti, R., Győry, F., Bacso, Z., Fésüs, L., & Kristóf, E. (2023). Extracellular thiamine concentration influences thermogenic competency of differentiating neck area-derived human adipocytes. Frontiers in nutrition, 10, 1207394. 10.3389/fnut.2023.1207394

Wang, C., Liang, X., Tao, C., Yao, X., Wang, Y., Wang, Y., & Li, K. (2017). Induction of copper and iron in acute cold-stimulated brown adipose tissues. Biochemical and biophysical research communications, 488(3), 496–500. 10.1016/j.bbrc.2017.05.073

Wang, T., Gao, Q., Yao, Y., Luo, G., Lv, T., Xu, G., Liu, M., Xu, J., Li, X., Sun, D., Cheng, Z., Wang, Y., Wu, C., Wang, R., Zou, J., & Yan, M. (2023). Causal relationship between obesity and iron deficiency anemia: a two-sample Mendelian randomization study. Frontiers in public health, 11, 1188246. 10.3389/fpubh.2023.1188246

Wenzel, B. J., Stults, H. B., & Mayer, J. (1962). Hypoferraemia in obese adolescents. *Lancet (London*, England), 2(7251), 327–328. 10.1016/s0140-6736(62)90110-1

Winnier, D. A., Fourcaudot, M., Norton, L., Abdul-Ghani, M. A., Hu, S. L., Farook, V. S., Coletta, D. K., Kumar, S., Puppala, S., Chittoor, G., Dyer, T. D., Arya, R., Carless, M., Lehman, D. M., Curran, J. E., Cromack, D. T., Tripathy, D., Blangero, J., Duggirala, R., Göring, H. H., … Jenkinson, C. P. (2015). Transcriptomic identification of ADH1B as a novel candidate gene for obesity and insulin resistance in human adipose tissue in Mexican Americans from the Veterans Administration Genetic Epidemiology Study (VAGES). PloS one, 10(4), e0119941. 10.1371/journal.pone.0119941

Wróblewski, K., & Kmiecik, S. (2024). Integrating AlphaFold pLDDT Scores into CABS-flex for enhanced protein flexibility simulations. Computational and structural biotechnology journal, 23, 4350–4356. 10.1016/j.csbj.2024.11.047

Yook, J. S., Thomas, S. S., Toney, A. M., You, M., Kim, Y. C., Liu, Z., Lee, J., & Chung, S. (2021). Dietary Iron Deficiency Modulates Adipocyte Iron Homeostasis, Adaptive Thermogenesis, and Obesity in C57BL/6 Mice. The Journal of nutrition, 151(10), 2967–2975. 10.1093/jn/nxab222

Yook, J. S., You, M., Kim, J., Toney, A. M., Fan, R., Puniya, B. L., Helikar, T., Vaulont, S., Deschemin, J. C., Okla, M., Xie, L., Ghosh, M. C., Rouault, T. A., Lee, J., & Chung, S. (2021). Essential role of systemic iron mobilization and redistribution for adaptive thermogenesis through HIF2-α/hepcidin axis. Proceedings of the National Academy of Sciences of the United States of America, 118(40), e2109186118. 10.1073/pnas.2109186118

Yook, J. S., You, M., Kim, Y., Zhou, M., Liu, Z., Kim, Y. C., Lee, J., & Chung, S. (2021). The thermogenic characteristics of adipocytes are dependent on the regulation of iron homeostasis. The Journal of biological chemistry, 296, 100452. 10.1016/j.jbc.2021.100452

Zhao, L., Zhang, X., Shen, Y., Fang, X., Wang, Y., & Wang, F. (2015). Obesity and iron deficiency: a quantitative meta-analysis. Obesity reviews : an official journal of the International Association for the Study of Obesity, 16(12), 1081–1093. 10.1111/obr.12323

Zhong, J., Zareifi, D., Weinbrenner, S., Hansen, M., Klingelhuber, F., Nono Nankam, P. A., Frendo-Cumbo, S., Bhalla, N., Cordeddu, L., de Castro Barbosa, T., Arner, P., Dahlman, I., Muniandy, M., Heinonen, S., Pietiläinen, K. H., Hoffmann, A., Ghosh, A., John, D., Tönjes, A., Ståhl, P. L., … Rydén, M. (2025). adiposetissue.org: A knowledge portal integrating clinical and experimental data from human adipose tissue. Cell metabolism, 37(3), 566–569. 10.1016/j.cmet.2025.01.012

