## Supplementary Material for "Iron uptake mediated by TFRC and secretion of transferrin types stimulate thermogenic activation in human adipocytes"

**Supplementary Materials**

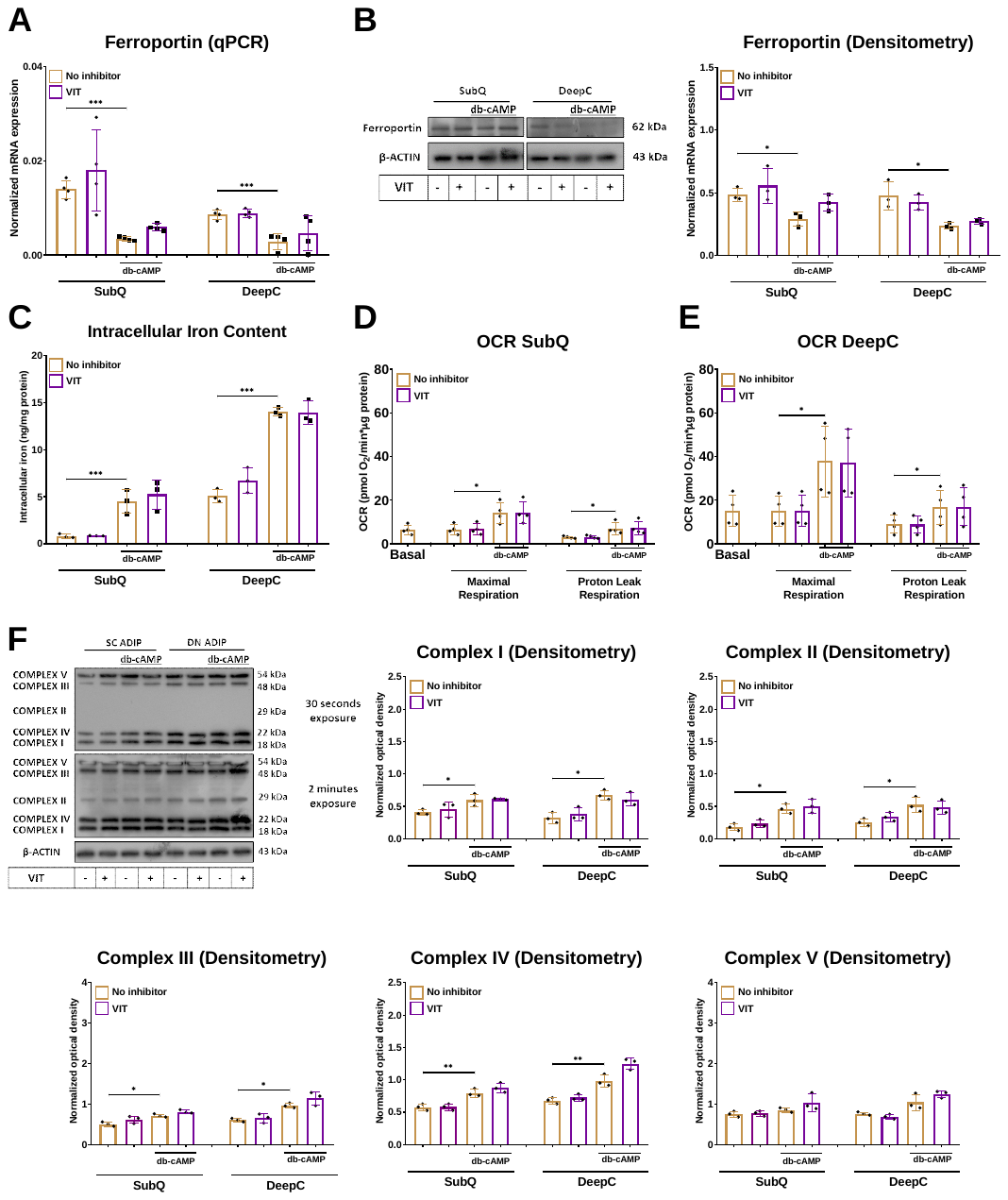

**G**
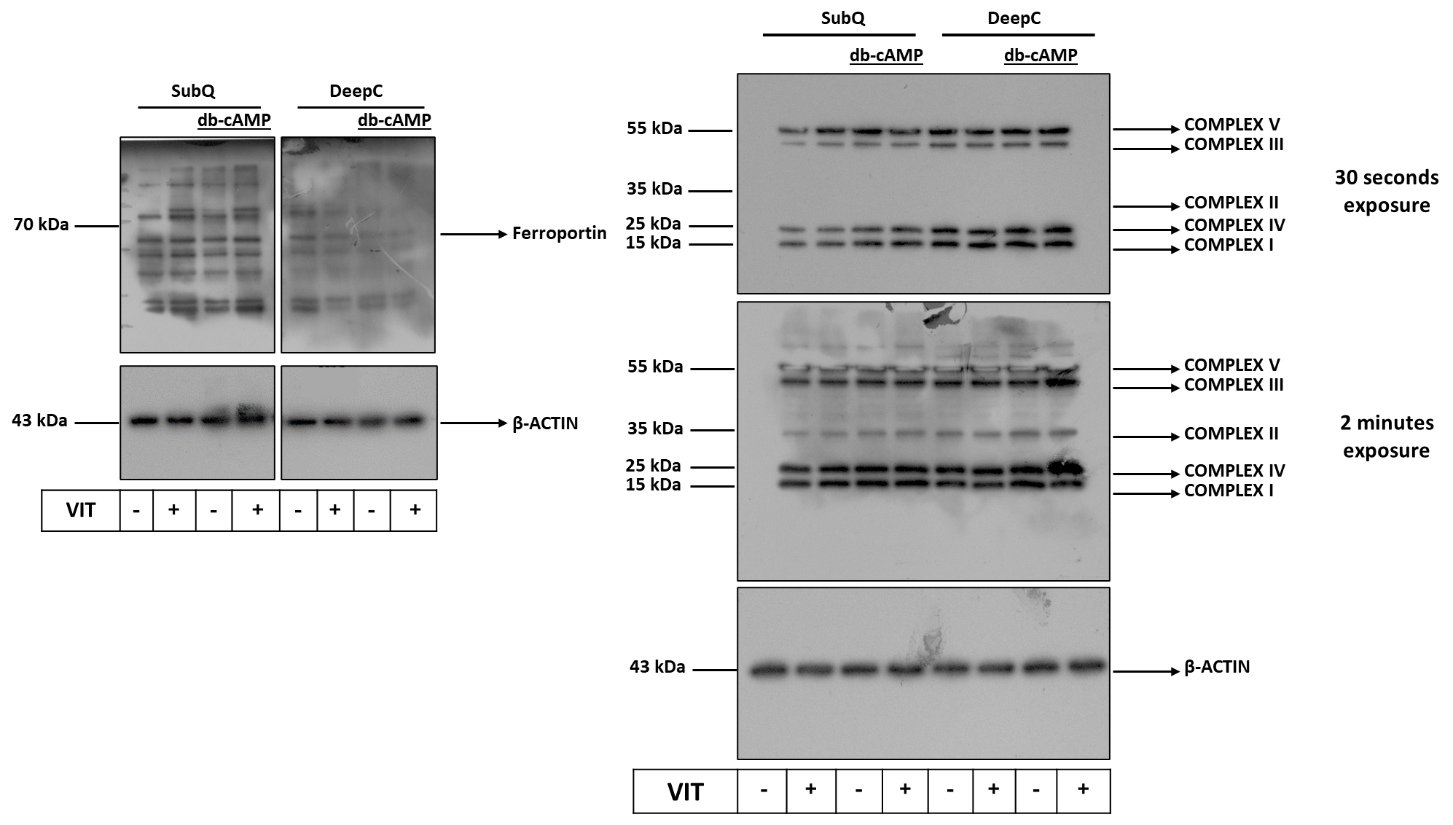

**Supplementary Figure 1. The effect of ferroportin inhibitor (****VIT-2763) on oxygen consumption and protein expression of mitochondrial complex subunits in human *ex vivo* differentiated subcutaneous (SubQ) and deep cervical (DeepC)-derived adipocytes.** SubQ and DeepC-derived adipocytes were treated with 500 µM dibutyryl (db)-cAMP, 200 nM VIT-2763, or combination of the two compounds for 10 hours. (A-B) The mRNA (A) and protein (B) expression of ferroportin (encoded by *SLC40A1*) detected by RT-qPCR (n=4) and western blot (n=3). (C) Intracellular iron content in SubQ and DeepC-derived adipocytes upon VIT-2763 treatment during adrenergic stimulation. (D-E) Basal, db-cAMP stimulated maximal, and proton leak oxygen consumption rate (OCR) in SubQ (D) and DeepC-derived (E) adipocytes were quantified by Seahorse extracellular flux analysis. (F) Protein expression of mitochondrial complex subunits detected by immunoblotting. (G) The original uncropped pictures of the full-length blots. Detailed information regarding the antibodies and working dilution are displayed in Supplementary Table 2. n=4, statistical analysis was performed by one-way ANOVA followed by Tukey’s *post hoc* test, *p<0.05, **p<0.01, and ***p<0.001.

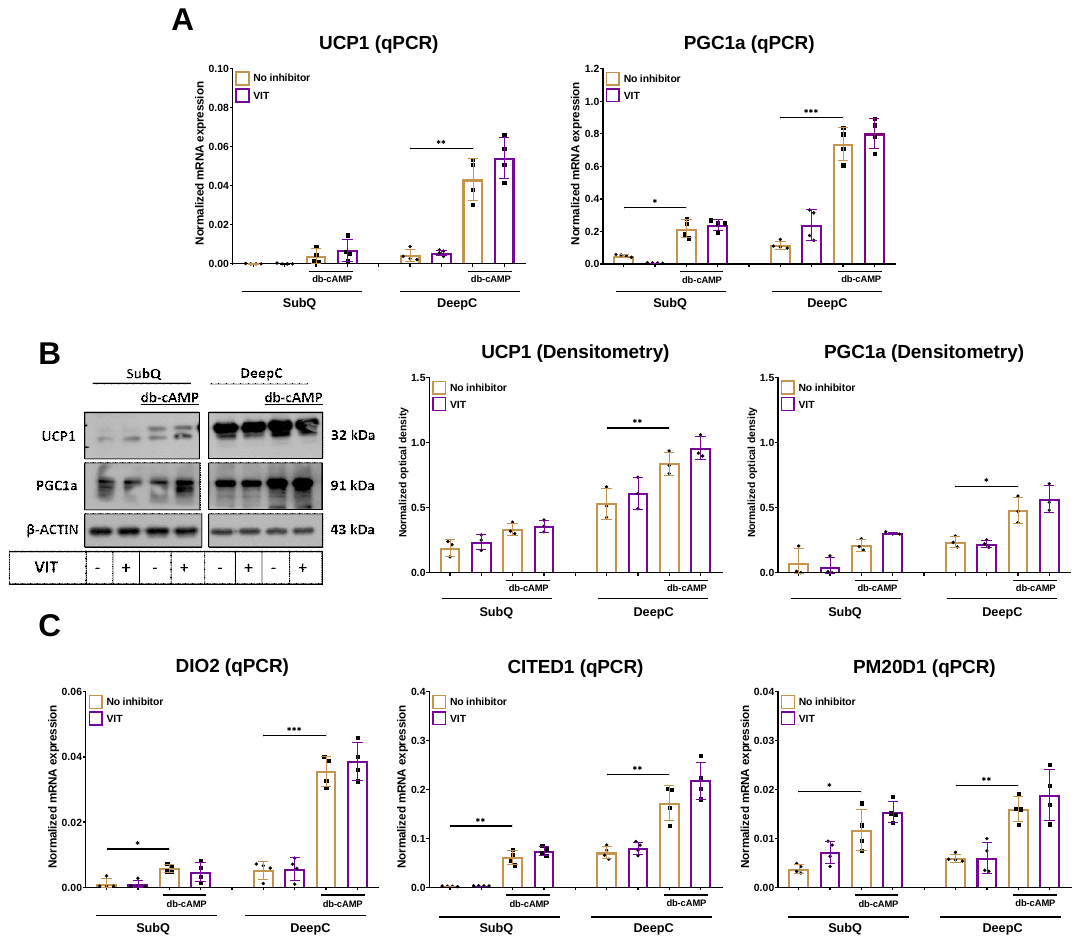

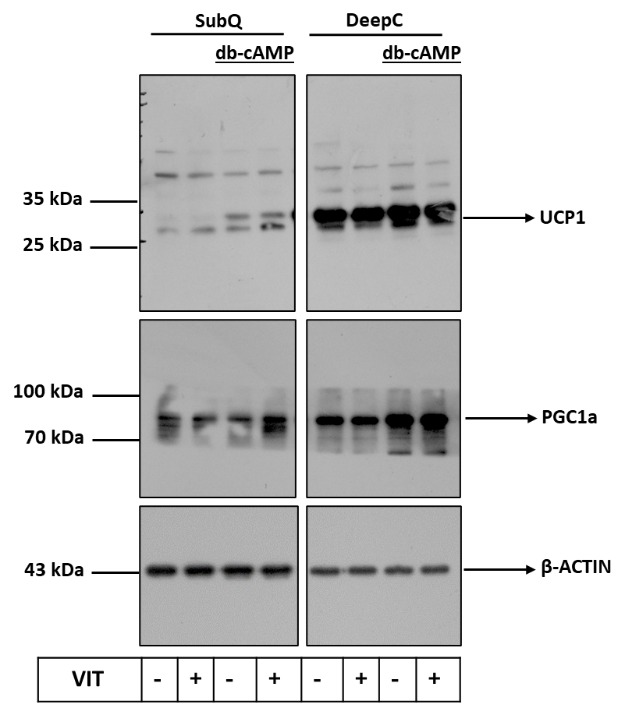
**D**

**Supplementary Figure 2. The effect of ferroportin inhibitor (VIT-2763) on the expression of thermogenic markers in human *ex vivo* differentiated subcutaneous (SubQ) and deep cervical (DeepC)-derived adipocytes.** Adipocytes were differentiated and treated as in Supplementary Figure 1. (A-B) mRNA (A) and protein (B) expression of UCP1 and PGC1a detected by RT-qPCR and immunoblotting, respectively. (C) mRNA expression of *DIO2*, *CITED1*, and *PM20D1* detected by RT-qPCR. (D) The original uncropped pictures of the full-length blots. Detailed information regarding the antibodies and working dilution are displayed in Supplementary Table 2. n=4, statistical analysis was performed by one-way ANOVA followed by Tukey’s *post hoc* test, *p<0.05, **p<0.01, and ***p<0.001.

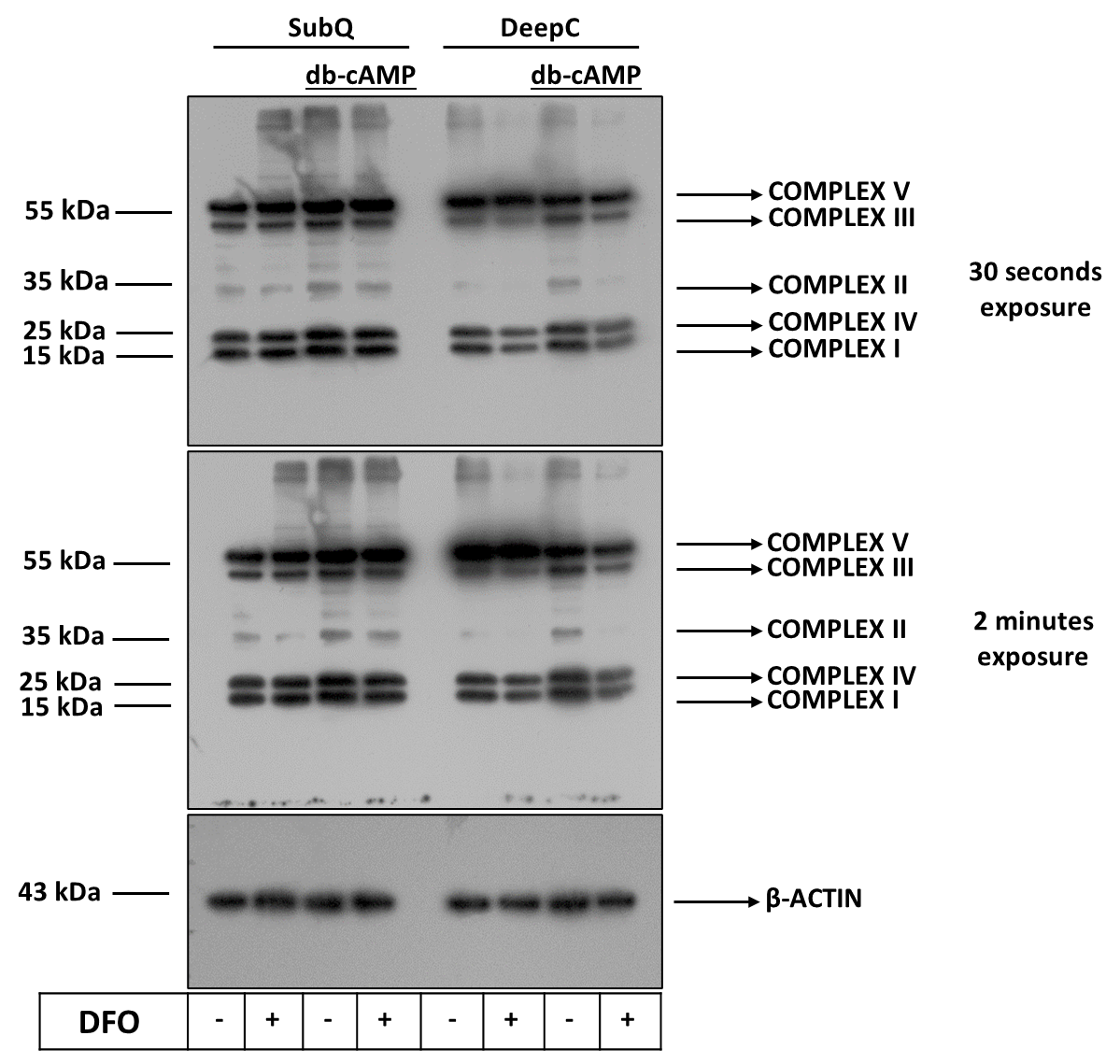

**Supplementary Figure 3. Uncropped images presented with molecular weight ladders for Figure 2D.** β-ACTIN was used as endogenous control. Detailed information regarding the antibodies and working dilution are displayed in Supplementary Table 2.

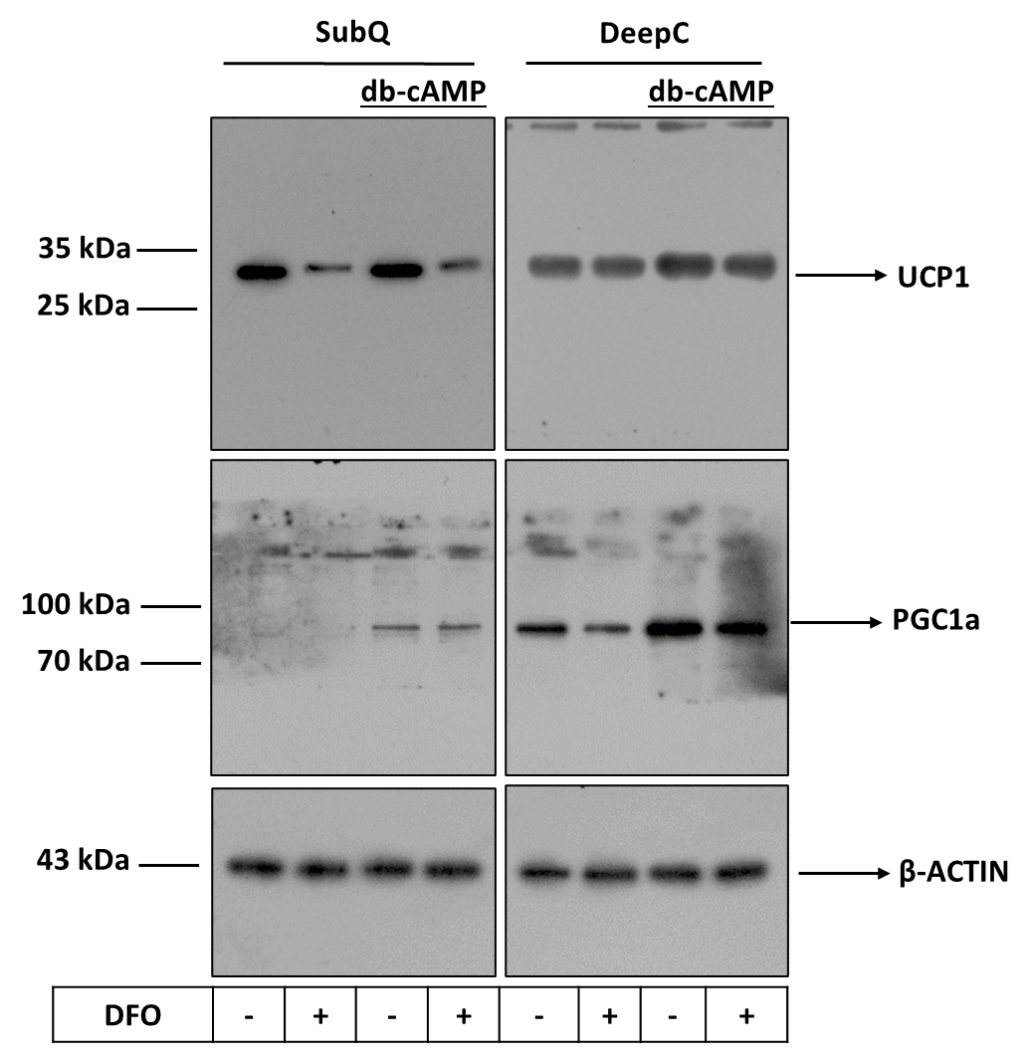

**Supplementary Figure 4. Uncropped images presented with molecular weight ladders for Figure 3B.** β-ACTIN was used as endogenous control. Detailed information regarding the antibodies and working dilution are displayed in Supplementary Table 2.

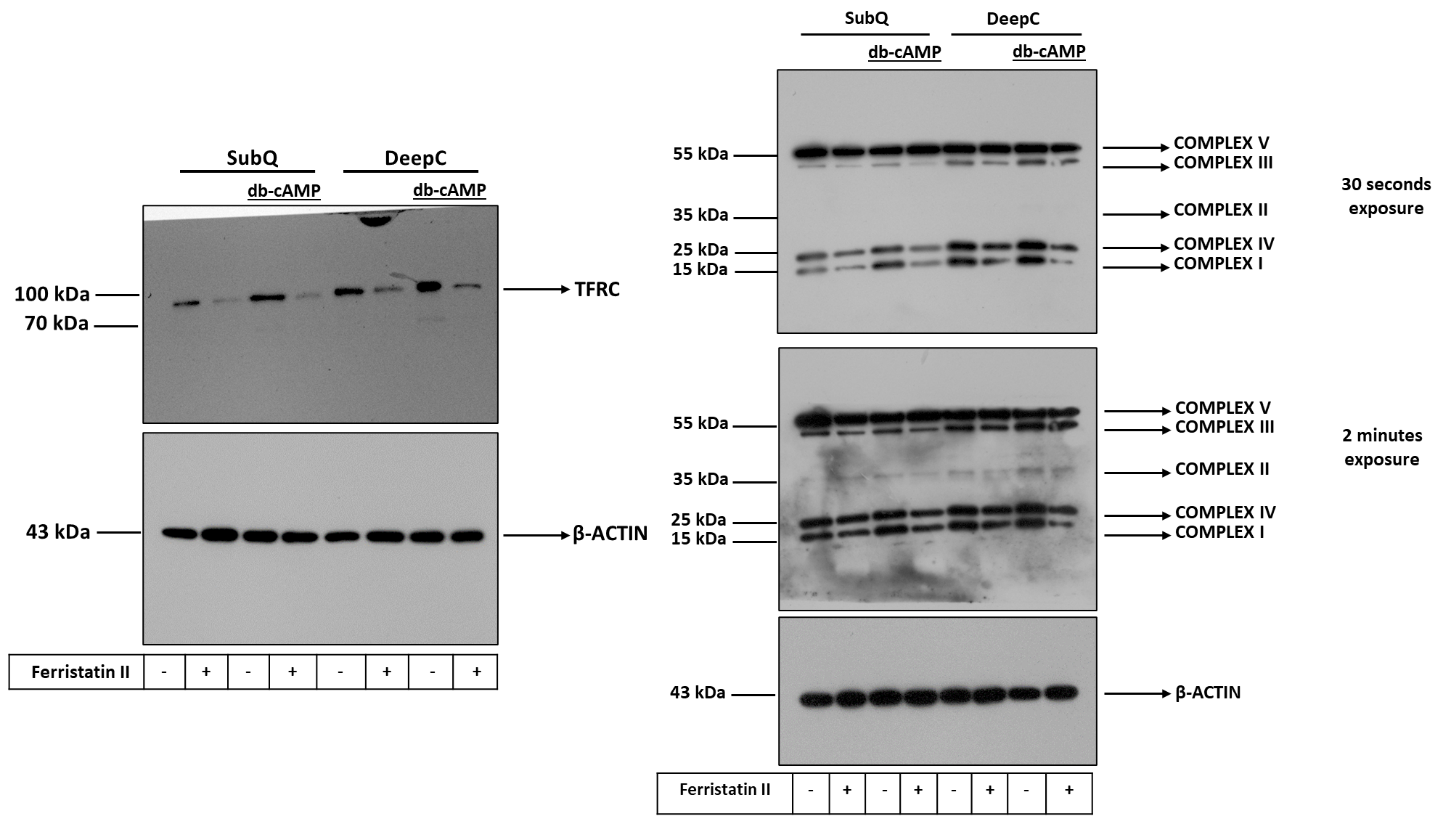

**Supplementary Figure 5. Uncropped images presented with molecular weight ladders for Figures 4B (left panel) and 4D (right panel).** β-ACTIN was used as endogenous control. Detailed information regarding the antibodies and working dilution are displayed in Supplementary Table 2.

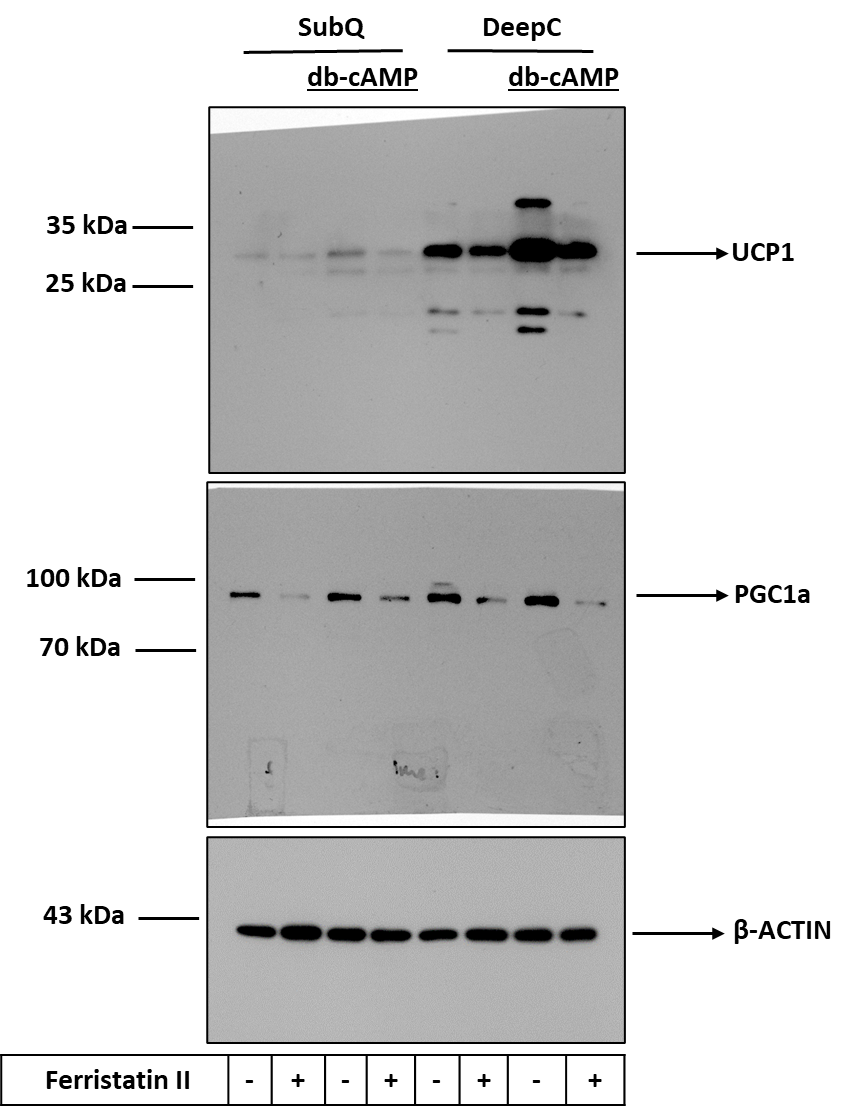

**Supplementary Figure 6. Uncropped images presented with molecular weight ladders for Figure 5B.** β-ACTIN was used as endogenous control. Detailed information regarding the antibodies and working dilution are displayed in Supplementary Table 2.

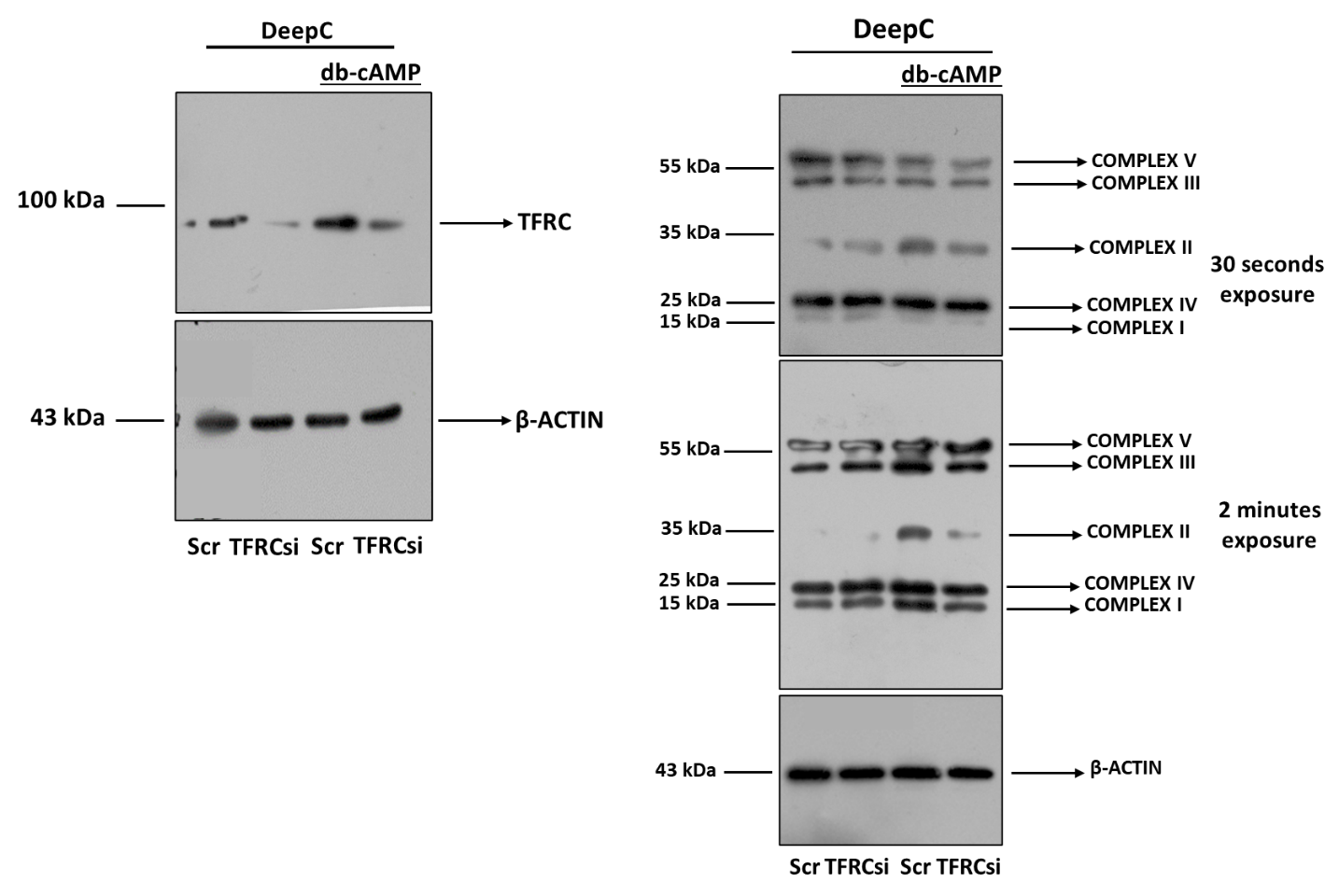

**Supplementary Figure 7. Uncropped images presented with molecular weight ladders for Figures 6B (left panel) and 6E (right panel).** β-ACTIN was used as endogenous control. Detailed information regarding the antibodies and working dilution are displayed in Supplementary Table 2.

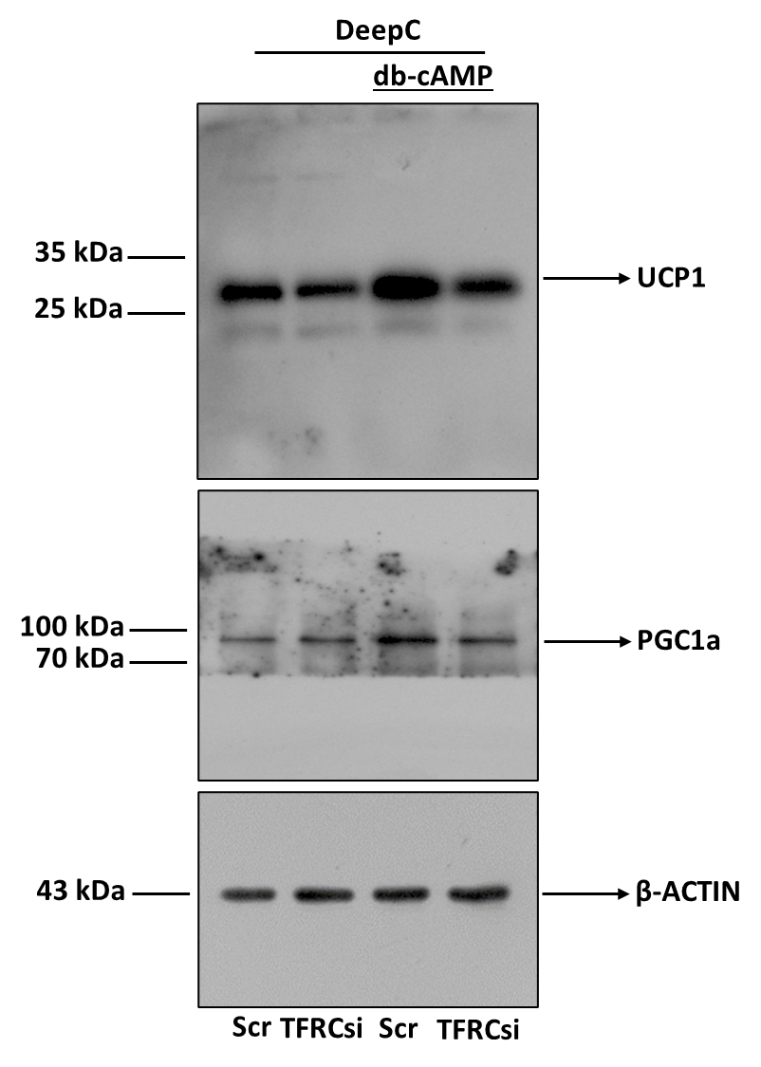

**Supplementary Figure 8. Uncropped images presented with molecular weight ladders for Figure 7B.** β-ACTIN was used as endogenous control. Detailed information regarding the antibodies and working dilution are displayed in Supplementary Table 2.

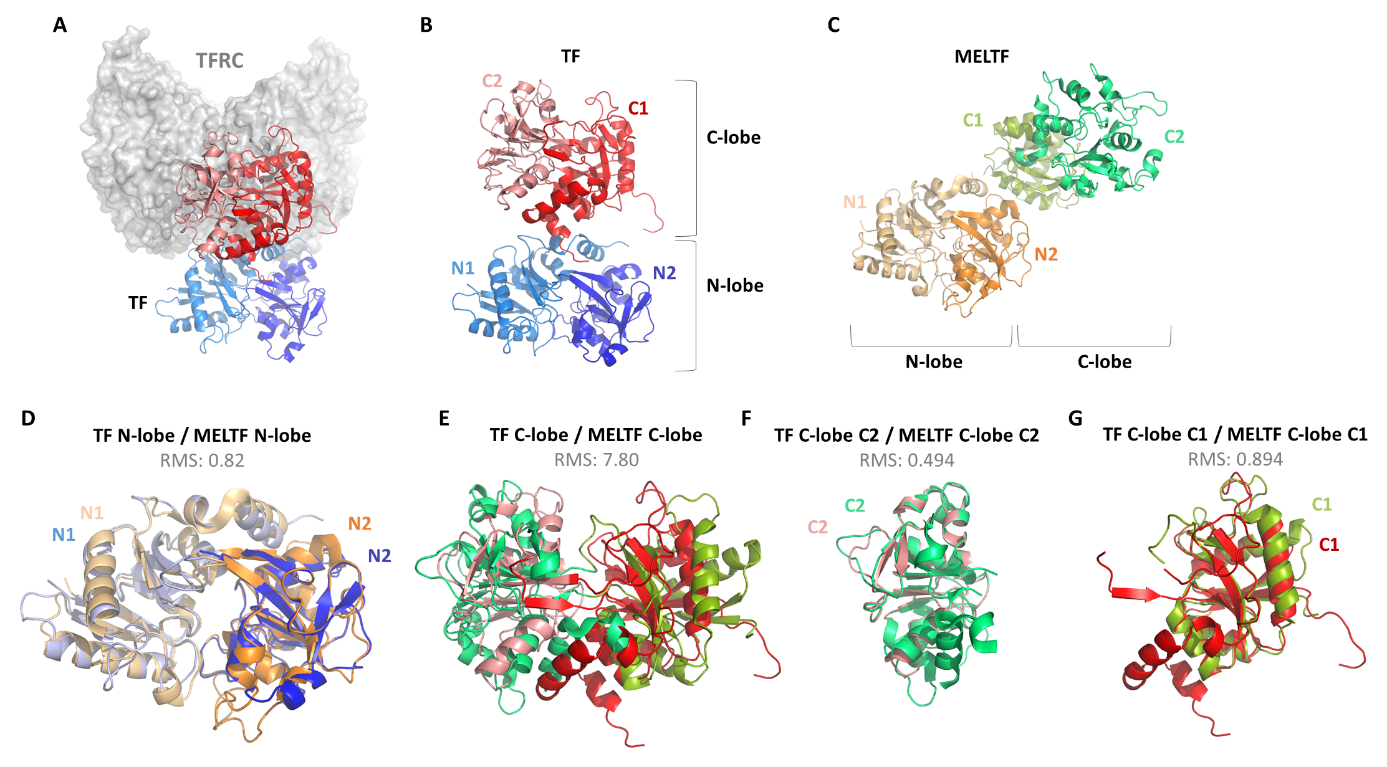

**Supplementary Figure 9. Comparison of N- and C-lobe domains and N1, N2, C1, and C2 subdomains of transferrin (TF) and melanotransferrin (MELTF).** (**A**) Structures of transferrin receptor (TFRC) complexed with TF (PDB ID: 1SUV). TFRC is grey, the N- and C-lobes of TF have blue and red colors, respectively. (**B**) Structure of TF (PDB ID: 1SUV). The subdomains of N-lobe (N1 and N2) and C-lobe (C1 and C2) are shown by different shades of blue and red colors, respectively. The N-lobe (1–331 residues) and the C-lobe (339–679) can be divided into two subdomains: N1 (1–92 and 247–331 residues) and N2 (93–246 residues), as well as C1 (339–425 and 573–679 residues) and C2 (426–572 residues) [Silva *et al.*, 2021]. (**C**) Structure of MELTF (PDB ID: 6XR0). The subdomains of N-lobe (N1 and N2) and C-lobe (C1 and C2) are shown by different shades of orange and green colors, respectively. (**D**) Alignment of N-lobe domains of TF and MELTF. (**E**) Alignment of C-lobe domains of TF and MELTF. (**F**) Alignment of C2 subdomains of TF and MELTF. (**G**) Alignment of C1 subdomains of TF and MELTF. RMS: Root Mean Square deviation.

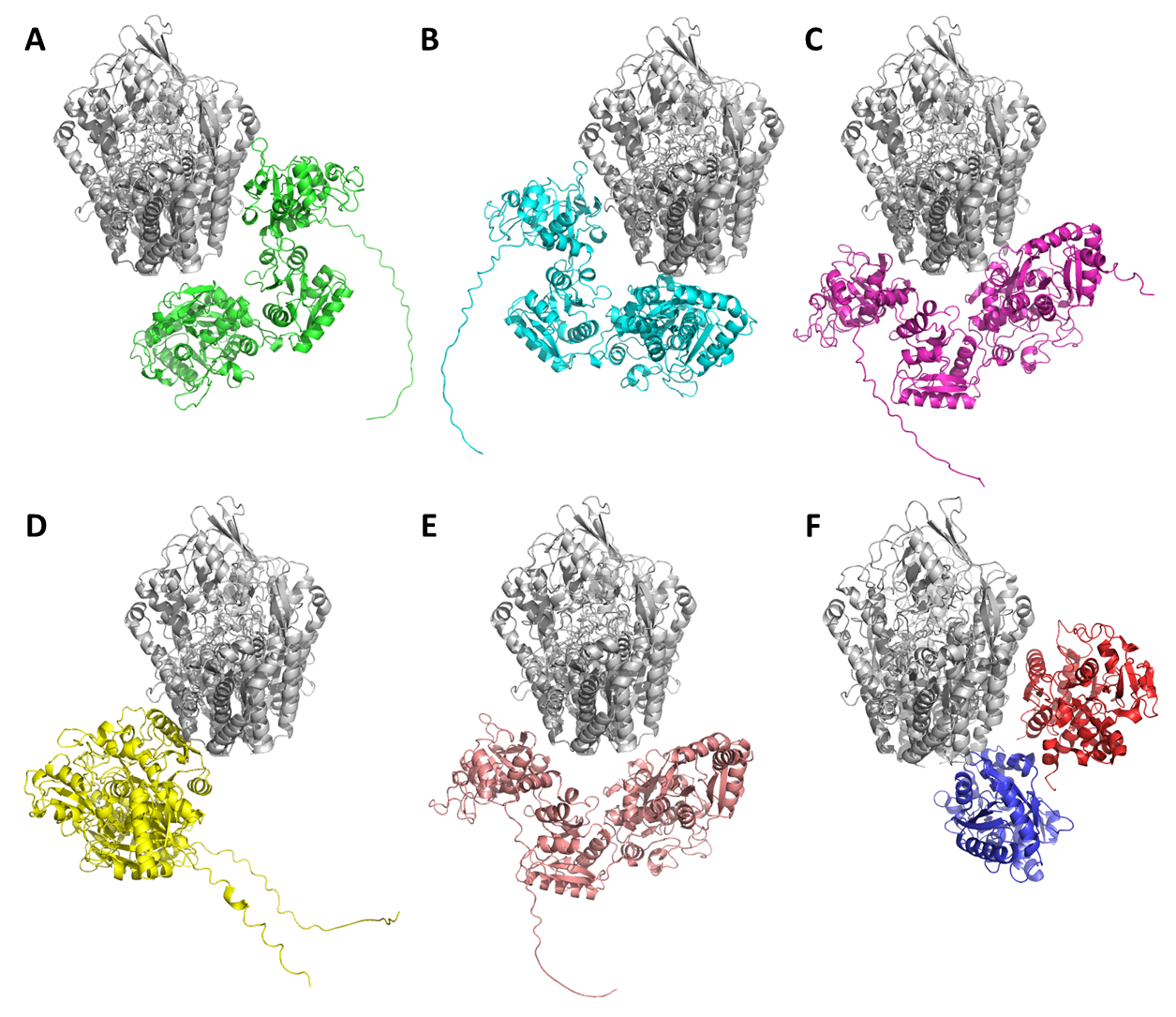

**Supplementary Figure 10. Possible binding modes of melanotransferrin (MELTF) to transferrin receptor (TFRC).** (**A-E**) Structures of TFRC complexed with MELTF were predicted and ranked by HADDOCK. Ribbon representations of the top five model structures are shown. TFRC is grey while MELTF has different colors in each case. MELTF is shown by green in the best model (A), the other top ranked models are shown in order (B-E). TFRC is shown in the same spatial orientation in every case. (**F**) The complex of TFRC and transferrin (TF) is shown for comparison, based on an electron microscopy structure (PDB ID: 1SUV) [Cheng et al., 2004]. The N- and C-lobes of TF have blue and red colors, respectively.

**Supplementary Table 1.** Gene primers and probes

| **GENES** | **ASSAY ID** |
| --- | --- |
| *ACTB* | Hs01060665**_**g1 |
| *CITED1* | Hs00918445_m1 |
| *DIO2* | Hs00255341_m1 |
| *GAPDH* | Hs99999905_m1 |
| *MELTF* | Hs00195551_m1 |
| *PM20D1* | Hs00399438_m1 |
| *PPARGC1A* | Hs01016719_m1 |
| *SLC40A1* | Hs00205888_m1 |
| *TF* | Hs00169070_m1 |
| *TFRC* | Hs00951083_m1 |
| *UCP1* | Hs00222453_m1 |

**Supplementary Table 2.** Antibodies used in immunoblotting

| **ANTIBODY** | **COMPANY** | **CATALOG NUMBER** | **DILUTION** |
| --- | --- | --- | --- |
| UCP1 | R&D Systems, Minneapolis, MN, USA | MAB6158 | 1:750 |
| TFRC | Santa Cruz, Dallas, TX, USA | 3B8 2A1 | 1:2000 |
| SLC40A1/Ferroportin | Invitrogen, Waltham, MA, USA | PA5-22993 | 1:2000 |
| PGC1α | Novus Biologicals, Centennial, CO, USA | NBP1-04676 | 1:1000 |
| Total OXPHOS | Abcam, Cambridge, UK | ab110411 | 1:1000 |
| ACTIN | Sigma-Aldrich, Munich, Germany | A2066 | 1:10000 |
| HRP-conjugated goat anti-rabbit IgG | Advansta, San Jose, CA, USA | R-05072-500 | 1:5000 |
| HRP-conjugated goat anti-mouse IgG | Advansta, San Jose, CA, USA | R-05071-500 | 1:5000 |

**Supplementary Table 3. Comparison of interactions of TF or MELTF with TFRC based on model complexes.** The values were obtained from the analysis of structural coordinates by using Generate module of PDBsum [Laskowski *et al.*, 2018].

| **Complex** | **TFRC + TF** | **TFRC + MELTF** |
| --- | --- | --- |
| **No. of interface residues** | 25 + 31 | 28 + 39 |
| **Interface area (Å)** | 1356 + 1315 | 1585 + 1412 |
| **Salt bridges** | 8 | 10 |
| **H-bonds** | 21 | 22 |
| **Non-bonded contacts** | 160 | 194 |
